# Parallel evolution of industrial melanism in the peppered moth: one locus, many alleles

**DOI:** 10.64898/2026.02.26.707722

**Authors:** Ivy Whiteford, Pascal Campagne, Arjen E. van’t Hof, Carl J. Yung, Michael Berenbrink, Shen Tian, Flora Todd, Jon Delf, Antónia Monteiro, František Marec, László Rákosy, Andrea J. Betancourt, Ilik J. Saccheri

## Abstract

The extent to which adaptation to environmental change occurs via single or multiple advantageous mutations remains an open question, which we examined by studying the spread of melanic forms of the peppered moth in Britain and continental Europe, in response to industrial coal pollution. In Britain, the darkest melanic form is due to the insertion of a transposable element (*carb-TE*) into a genomic region called *ivory*. Here, we characterize the spread of melanic forms in continental Europe from historical records and uncover the genetic basis of European melanism using genomic analyses of modern and museum specimens. We show that European melanism is also associated with variants at *ivory*, but with multiple alleles featuring structural variants, including several transposable elements, though not *carb-TE*. The primary central European melanic allele is a genetically dominant 805 base pair deletion (*sollichau*). Interestingly, melanic individuals with either *sollichau* (deletion) or *carb-TE* (insertion) alleles show elevated expression of *ivory* and its effector micro-RNA (*mir-193*) compared to non-melanic (*typica*) individuals, suggesting that contrasting structural variants in *ivory* have similar regulatory effects. A functional role for *sollichau* was corroborated, serendipitously, by its presence in a *typica* individual which also contains a linked deletion of *mir-193*, which is predicted to cancel the melanizing effect of the *sollichau* deletion. Our results support the idea that there can be many genetic origins of the same adaptive trait within a species, particularly one with a large effective population size and heterogeneous natural habitat, and when many mutations can give rise to the same phenotype.

## Introduction

By the mid 19th century, the emergence of new industries fueled by coal combustion resulted in the accumulation of environmentally harmful particulate and volatile emissions, e.g., black carbon (soot), sulfur dioxide (SO_2_), and carbon dioxide (CO_2_). Britain was at the forefront of this dramatic socioecological transformation, later paralleled in parts of Europe, North America and Asia. In this post-industrial epoch – the Anthropocene – human-associated environmental change drives mass extinctions, extensive range shifts, and rapid adaptation ^1^.

In a textbook case of adaptation to pollution ^2^ a melanic variation (‘*carbonaria’*) of the peppered moth (*Biston betularia*), first documented near Manchester in 1848, became the predominant form of the species within decades. The rise of *carbonaria* was attributed to a single *de novo* allele originating shortly before its spread, driven by strong positive selection ^3,4^ – an interpretation later corroborated by the identification of a single causal mutation with an inferred date of origin in the early 1800s ^5^. Surprisingly, *carbonaria* was due to a transposable element insertion (*carb*-TE), in the region of two overlapping genes, *cortex* and *ivory*, a hotspot for wing-pattern evolution throughout Lepidoptera ^6^. In particular, structural variation around *ivory*, a long non-coding RNA (lncRNA), has been linked to multiple polymorphic wing phenotypes ^7–9^. Furthermore, deletion of an associated microRNA, *mir-193*, results in a loss of wing coloration in three butterflies ^9^ and a moth ^10^.

Shortly after the first reports of *carbonaria* in Britain, similar melanic *B. betularia* were reported from separate locations in northwestern Europe, in parallel with the onset of coal pollution (Breda, Netherlands in 1867; Halle, central Germany in 1874, and Bremen, northern Germany in 1879; Figure 1a and b, Table S1). The spread on the continent appeared to follow the progress of coal mining and industrialization from western to eastern regions. This led to debate both about the nature of the polymorphism - heritable or environmentally induced - and the source of the black variant (Table S1). In particular, Hasebroek ^11^ and Ule ^12,13^ recognized that the timing and spatial pattern of early records of the black morph on the continent suggested multiple local sources (Figure 1a), rather than a long distance migration event from Britain.

**Figure 1.**
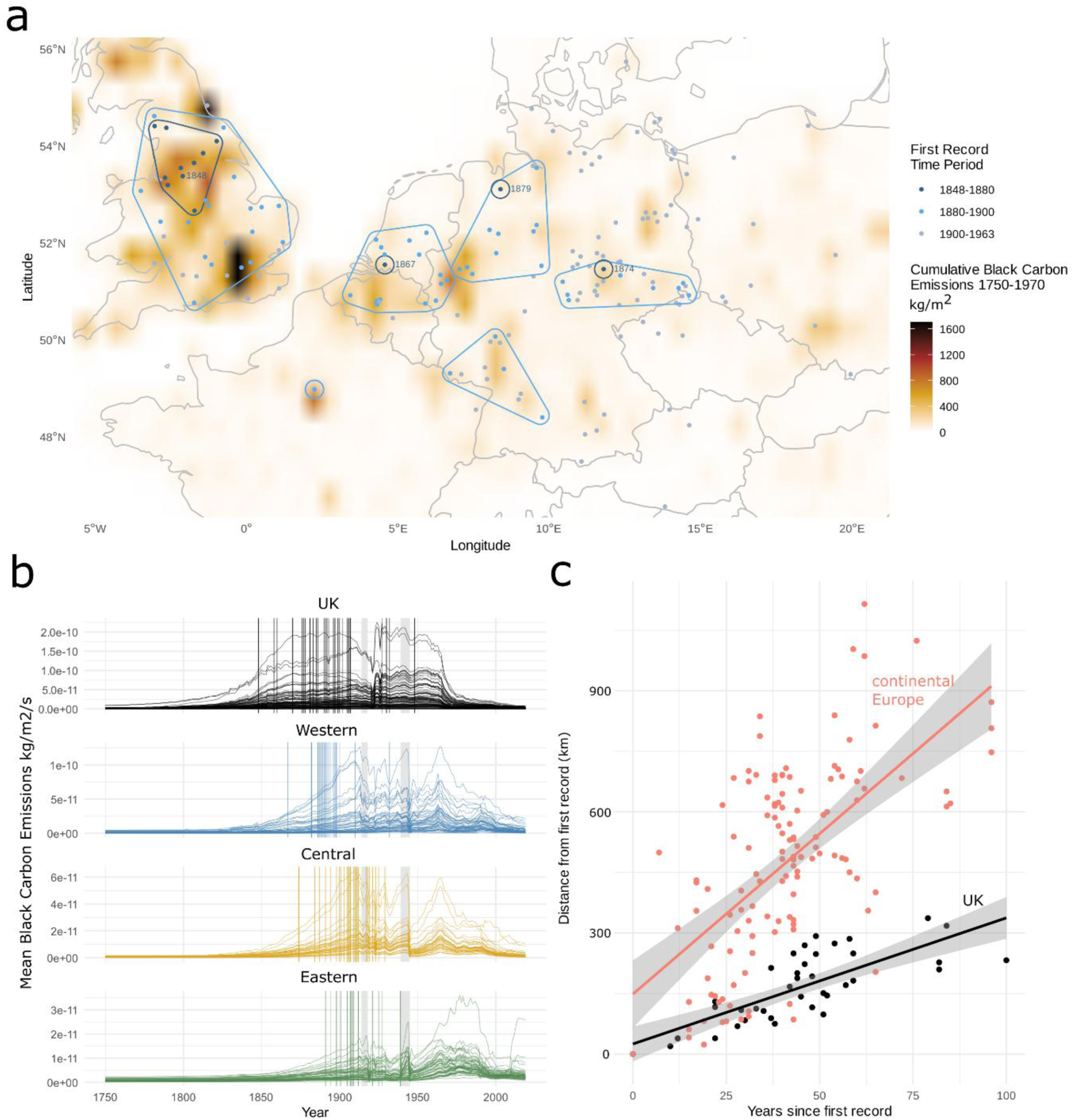
The spread of industrialisation and melanic variation in the peppered moth. (**a**) A map of mainland Britain and northwest continental Europe showing the estimated cumulative black carbon particulates emitted from 1750-1970 and published first records of melanic *Biston betularia* (Video S1 shows animation of progress of SO_2_ concentration, a correlate of soot pollution). First record locations are clustered by distance and outlined with colours indicating time periods; for the earliest time window (1848-1880), the year of the first record within each cluster is shown. (**b**) Black carbon emissions throughout the Anthropocene, divided into four study areas comprising the UK (mainland Britain), a Western study region (an area across the west of Germany, Belgium, Netherlands), a Central study region (an area across the east of Germany) and an Eastern study region (an area across Czech Republic and southwestern Poland). Curves indicate measurements from individual grid squares. Vertical annotation lines show the timing of local first records of melanic *B. betularia*. Grey shading indicates the two World Wars. (**c**) Comparison of the implied rate of spread (regression slope) of melanic *B. betularia* in Britain (black, slope=3.12, R^2^=0.57) and continental Europe (red, slope=7.95, R^2^=0.36) under the assumption of single local origins in both landmasses. Lines indicate a linear regression, with shaded areas indicating 0.95 confidence intervals.

We have used an expanded set of first records to estimate the rate of spread of the black form, separately for mainland Britain and continental Europe, under the assumption of a single genetic origin within each landmass (Figure 1c). Consistent with previous interpretations, our analysis shows that the black form appeared to spread much more rapidly in continental Europe than Britain, 8 vs 3 km/year on average, with implausibly high rates required to explain the many outliers, unless multiple origins are considered (Figure S1). In fact, melanism is known to have arisen multiple times, in the form of partially melanic *insularia* ^14^ and North American *swettaria* morphs of *B. betularia* ^15,16^, and in many other Lepidoptera species ^17^. Despite independent origins, melanism may involve similar genes, with the *ivory/cortex* region implicated in other geometrid moths ^10,18^.

The gradual introduction of pollution controls starting in the 1950s produced a decrease in soot and SO_2_; in Britain, steep declines in melanic *B. betularia* followed, with the *carbonaria* form now nearly extinct ^5,19,20^. In the Netherlands, the continental European country with the best records, there were parallel declines ^21^. Notably, in Eastern Bloc countries, the 1973 oil price crisis led to greater dependency on coal ^22^, and the rigidity of the economic system meant that pollution controls were instituted only after the major political restructuring events of the 1990s (Figure S2).

Here, we study industrial melanism of *B. betularia* from northwest and central Europe. We confirm that, as in Britain, melanic records for this species were strongly associated with industrial pollution. The genetic basis of European melanic forms is architecturally simple, mapping to the *ivory/cortex* region as in British moths. However, unlike in Britain, the mutational basis cannot be assigned to a single origin, probably reflecting a larger effective population size on the continent. This work illustrates that parallel evolutionary responses to environmental change among, and even within, populations of the same species may be common where mutations of large effect arise frequently, and perhaps persist at low frequencies under heterogeneous ecological conditions.

## Results

### High frequency of melanic morphs in eastern Europe is associated with historical coal pollution

To describe melanism in wild-caught *B. betularia* across western and central Europe, which vary semi-continuously in the proportion of light and dark scales covering the upper wing and body surfaces (Figure 2a), we use the seven category scale of Cook and Muggleton ^23^, with A-B considered *typica*, C-F *insularia*, and G fully melanic. Crosses show that these phenotypes form a dominance series, with darker morphs dominant to lighter ones, and the G morph fully dominant ^14^. As *insularia* alleles show partial dominance and plasticity (Figure S3) ^14^, wild-caught material cannot be assigned to definite allele classes with certainty. Exploratory sampling in continental Europe between 2000 and 2012 directed us to concentrate our sampling efforts on three broad regions: eastern Germany (‘Central’ region), southwestern Poland and the Czech Republic (‘Eastern’ region), and western Germany and the Netherlands (‘Western’ region; Figure 2b; Table S2; Data S1).

**Figure 2.**
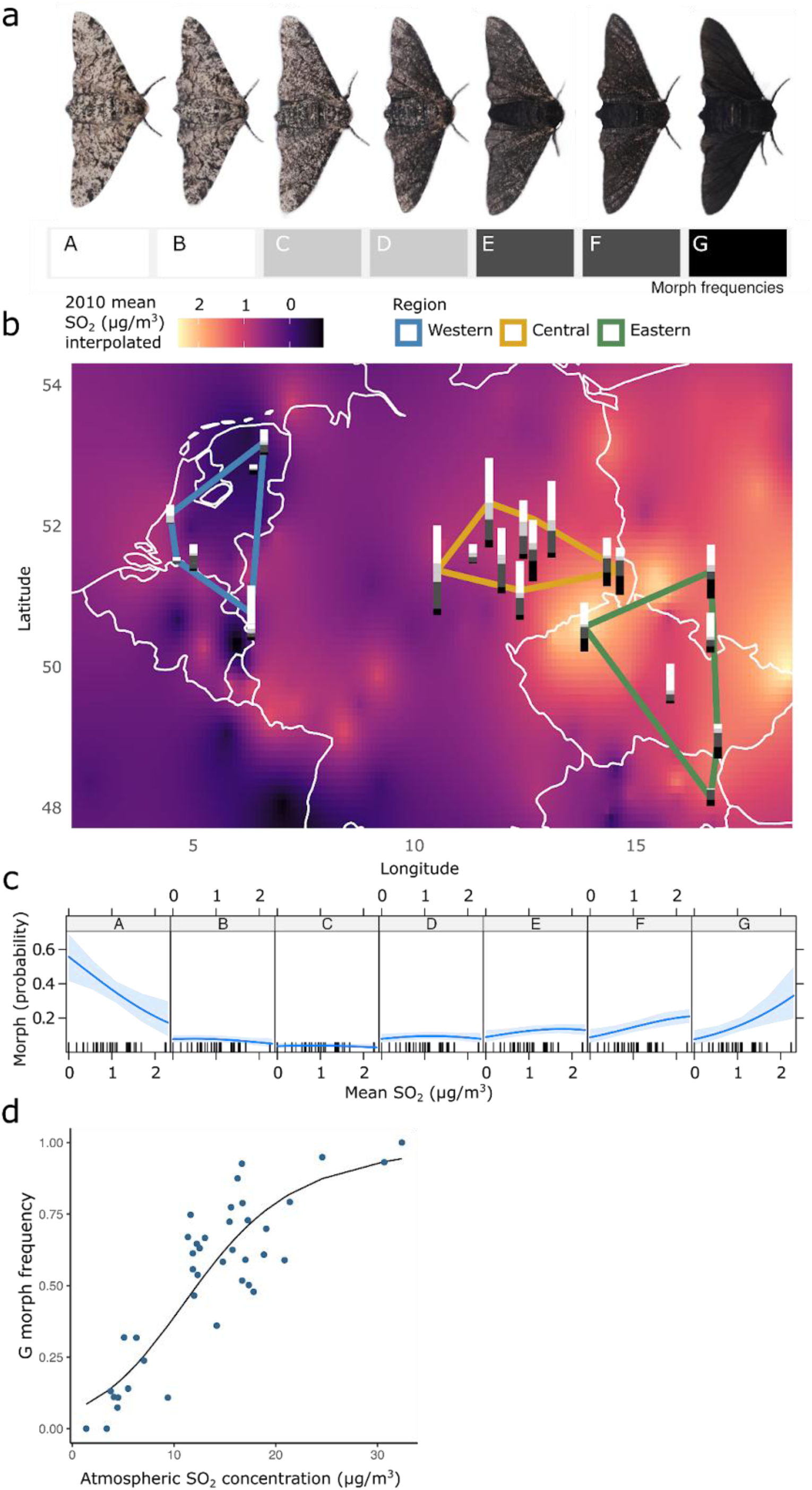
The melanic phenotype of *Biston betularia* is variable and associated with coal pollution. (**a**) Examples of the range of phenotypes found across European sampling locations, categorized into 7 morphs (A-G). (**b**) Contemporary (2013/14) frequencies of morphs observed at different sampling sites across ‘Western’ (blue), ‘Central’ (yellow) and ‘Eastern’ (green) regions, scaled to sample size (range = 5-70). An interpolated surface of historic SO_2_ concentration is overlain (AirBase 2010 data, rural stations, daytime sampling). (**c**) The effect sizes of interpolated atmospheric [SO_2_] on morph sampling probability when modelled with ordinal logistic regression. (**d**) Relationship between the frequency of the darkest (G) morph and [SO_2_] in the former Czechoslovakia, based on a sample of 4,939 moths collected between 1976-1982 ^24^ and 1995 [SO_2_] from AirBase interpolation.

In 2013/14, we sampled and scored the phenotype of 525 *B. betularia* from these three regions (Central: 321, Eastern: 121, Western: 83). We found that G-morph, full melanics, were at higher frequency in Central and Eastern Europe (0.15 and 0.25, respectively) than in the West (0.08; Table S2), consistent with a 30-year/moth generation (1960s vs 1990s) delay to the introduction of soot pollution controls in the former Eastern Bloc regions. Overall, the frequencies of the extreme morphs (A and G), but not the intermediate morphs, were weakly correlated with contemporary (2010) SO_2_ concentration (Figure 2c), reflecting erosion of historically steep SO_2_ gradients, paralleling changes observed in northwest England ^20^. A much stronger correlation with 1995 [SO_2_] is shown by the reanalysis of historical data on G morphs from the former Czechoslovakia during a period of high coal pollution in the 1970s and 80s ^24^, with melanism frequencies exceeding 90% at the highest [SO_2_] (Figure 2d).

### All melanic morphs map to a single locus containing the long non-coding RNA ivory

PCR-based genotyping of a small sample of melanic males wild-caught in the Netherlands (2007/08) and eastern Germany (2011) established that none of these individuals carried the UK *carbonaria*-TE (Figure S4). Linkage mapping using families derived from two of these males, Sollichau-G and Ramstedt-F (Figure S4), showed that melanism maps to the *ivory/cortex* region in both cases (Figure S5, Table S3). Genotyping of 335 individuals from the 2013/14 sample at 32 polymorphic markers spanning 124kb around the *cortex-ivory* region (Table S4) revealed that F and G morphs harboured several core haplotypes in this region, but the markers were too sparse to determine whether this reflected different functional alleles or the same alleles on recombined haplotypes.

To further resolve the genetic basis of melanism in these populations, we generated short-read sequence data for 637 individuals, wild-caught from 2000-2014 (*n* full melanic = 108; *n insularia* = 230; *n typica* = 299) and 17 laboratory-reared individuals, as well as 137 melanic (F and G) individuals from museum collections, aimed at the early 20^th^ century phase of melanism spread in Western and Central regions. To identify all types of genetic variants that might be associated with melanism, including large structural variants, we constructed a panel of complete reference haplotypes, informed by a preliminary genotyping approach (Figure S6). We sequenced a subset of eight melanic individuals using PacBio HiFi reads, assembled them into 16 haplotypes, aligned them to the British reference genome (accession GCA_905404145.2), and resolved these into a graph structure in which variation is stored as alternative paths along a common reference (hereafter referred to as a pangenome) ^25^. Six of these individuals were wild-caught and have at least one haplotype containing a melanic allele. Two were the offspring of crosses between melanic and *typica* individuals from the mapping families described above; for these individuals, the haplotype inherited from the melanic parent was identified using Illumina reads. Using the 6.6M variants discovered in the pangenome graph, we then genotyped 800 samples sequenced with short-reads, from across the study transect and museum specimens, using a k-mer based genome inference method, resulting in 1.9M loci (pangenie GQ>200, max-missingness 95%). This k-mer genotyping approach is limited to genetic variants present on the 16 assembled haplotypes and the reference, but allows efficient genotyping from short-read data, including the museum sample, both for SNPs and structural variants.

We contrasted the genotypes of each melanic morph category (D, E, F, G) with non-melanic morphs combined (A and B). We observe a clear singular outlier peak of genetic differentiation on chromosome 17 (corresponding to Merian element 7) ^26,27^ for the uniformly melanic G morph, with weaker peaks at the same locus for the remaining intermediate morphs (Figure 3a). Inspection of these peaks shows that the strength of the association peak is roughly proportional to the degree of melanism, such that the strongest peak is for the AB vs G contrast, closely followed by AB vs F, somewhat weaker for the AB vs E contrast, and weakest for AB vs D. Within the *ivory* region (Figure 3b), this differentiation spans ∼200kb from a region upstream of the conserved *ivory* promoter, containing *cortex* and orthologs of putative cis-regulatory elements of *ivory* implicated in wing pattern variation of the butterfly *Junonia coenia* ^7^, to ∼40kb before miRNAs 193 and 2788.

**Figure 3.**
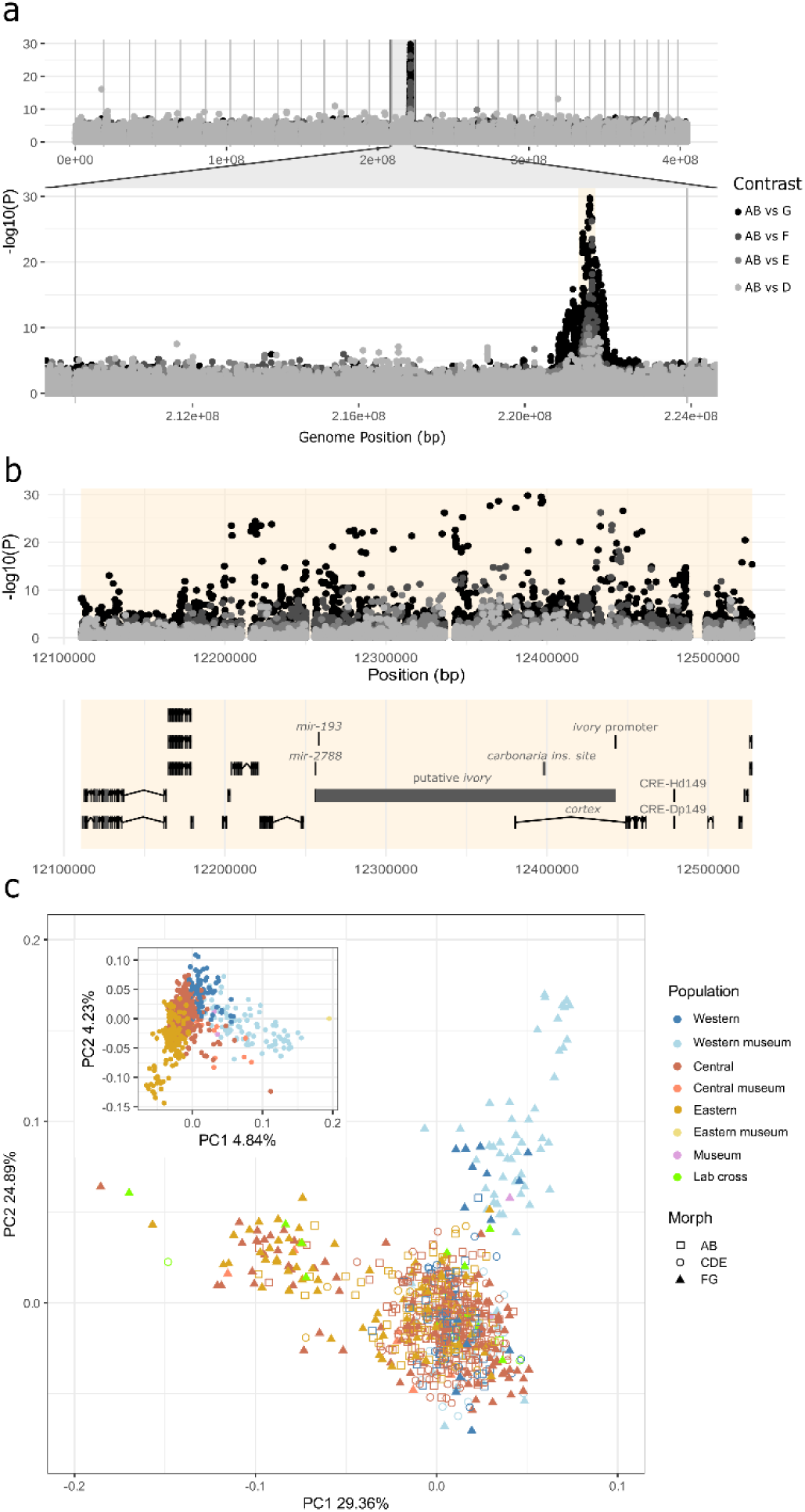
Melanic morphs are associated with genotypes in the *ivory* region. (**a**) Genome-wide and outlier chromosome genotype associations for each individual melanic category (D-G) vs the *typica* (AB) sample (*ivory* region highlighted in beige). Individual chromosomes are delineated by vertical grey lines. (**b**) *ivory* region genotype associations (including museum samples) for each individual melanic category (D-G) vs the *typica* (AB) sample and annotation: historical candidate gene *cortex* and functional UK *carbonaria*-TE insertion polymorphism; *ivory* precursor lncRNA and associated miRNA sequences; and CREs, predicted from sequence similarity ^7,9^ and transcriptomic evidence (SRR1021599 & ERR10123640). (**c**) Principal component analysis of SNP genotyping data across the *ivory* region (main plot) and the whole genome (inset plot). Museum specimens labelled in pink originate outside of the defined study areas.

### Peppered moth melanism is defined by several alleles in the *ivory* region

An alternative approach to partitioning the SNP variation in the *ivory* region using principal component analysis (Figure 3c), suggests two separate geographic clusters of the darkest melanic forms (mostly G) and a third large cluster containing a varied mix of melanics and non-melanics. One F/G cluster is composed of Western samples, mostly (but not exclusively) museum, while the other is composed of Central and Eastern samples, mostly modern with only two museum samples. The high proportion of museum melanics in the Western sample, compared to the other two study areas, is due to the low frequencies of melanics in the modern Western region, and smaller museum sample for Central and Eastern regions. These clusters do not reflect population structure – whole genome PCA (Figure 3c inset) shows weak spatial genetic structuring across the three study areas (on PC2), consistent with large effective population sizes (Figure S7), weak population structure (Figure S8), and high rates of dispersal^20^. The separation of whole genome museum samples on PC1 is likely due to technical differences in library preparation (single- vs double-stranded DNA), and does not affect the clustering pattern observed for the *ivory/cortex* region.

### Many melanism-associated haplotypes in continental Europe

We explored the type and co-occurrence of loci showing strong associations with melanism (Figure 4a; *p* < 0.01 for AB vs D, E, and F contrasts, and *p* < 1e-08 for AB vs G contrast). A number of highly associated polymorphisms appear to be structural variants, in some cases located close to the region of the *carbonaria*-TE insertion identified in British melanic G forms, and in other cases further downstream of the *ivory* lncRNA. Analysis of multilocus genotypes (MLG) indicates the presence of multiple distinct melanic haplotypes (Figure 4b). The darkest phenotype, the G category, is comprised of individuals with several MLGs, two including variants with significant association with melanism. MLG groups in individuals with intermediate, *insularia* phenotypes, show more phenotypic variation, with similar MLGs appearing in multiple morph categories. This effect can also be observed within controlled families (Figure S3), and is consistent with partial dominance and/or plasticity of these types. Due to the strong linkage disequilibrium between markers and greater rates of missing data observed in indel genotyping, it is difficult to untangle the causative polymorphism from linked neutral variants in these core haplotypes.

**Figure 4.**
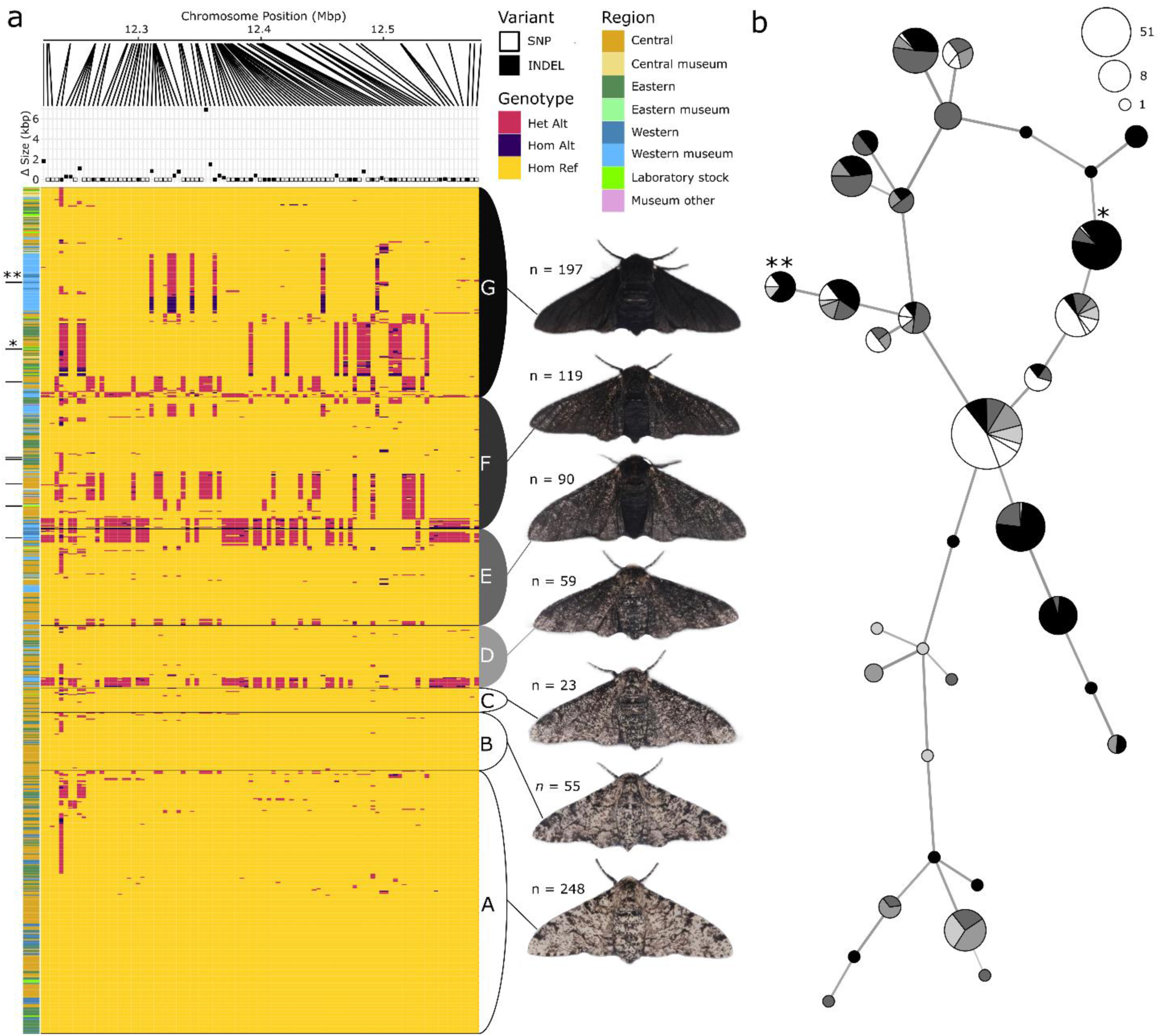
Melanic morph associated genotypes occur in distinct multilocus genotype groups which can produce varied melanic phenotypes. (**a**) A subset of markers independently associated with one or more melanic phenotypes are shown across the *ivory* region. The size of the polymorphism at each locus (SNP=0; indels of varying size) accompanies an indication of their physical position on the M7 chromosome. Genotypes for each sampled individual at these markers are shown in rows, grouped by phenotype category (A-G). To the left of the genotypes, black lines indicate individuals assembled with HiFi reads, and the coloured lines indicate a sample’s source population. (**b**) A contracted (threshold=7) multilocus genotype network of the same individuals and melanism associated loci. Single and double asterisk identify the primary G-associated genotype in the Eastern and Western regions, respectively.

### Diverse structural variants, including TEs, are associated with melanism

We were particularly interested in the role of large structural variants in the *ivory/cortex* region, as a large TE insertion is implicated as the causal mutation in the UK *carbonaria* allele. In our 16 long-read haplotypes, there were seven new large insertions or deletions relative to the reference in this region (Figure 5a). The variants were due to diverse causes, with five TE insertions (four of which are independent) from potentially active TE families (Table S5): one is LINE/L2 element (G1_2013_18.2); two from different LTR/Pao families (G6_2013_44.2 and DW_2013_04.1); and two are alleles of the same PiggyBAC TEs insertion (HA_2013_18.2 and ZU_2013_05.1). Two additional sequences appear to have arisen through tandem-duplication (DW_2013_04.2) and deletion (*sollichau*). The prevalence of structural variants in the *ivory/cortex* region is typical for a region of this size (Figure 5b); we also find that the rate of recombination is similar to that of other chromosomes. However, without precise knowledge of regulatory element boundaries and a genome-wide map of comparable regulatory sequences, it is difficult to refine this analysis further to address whether regulatory elements controlling *ivory* expression are enriched for structural variation.

**Figure 5.**
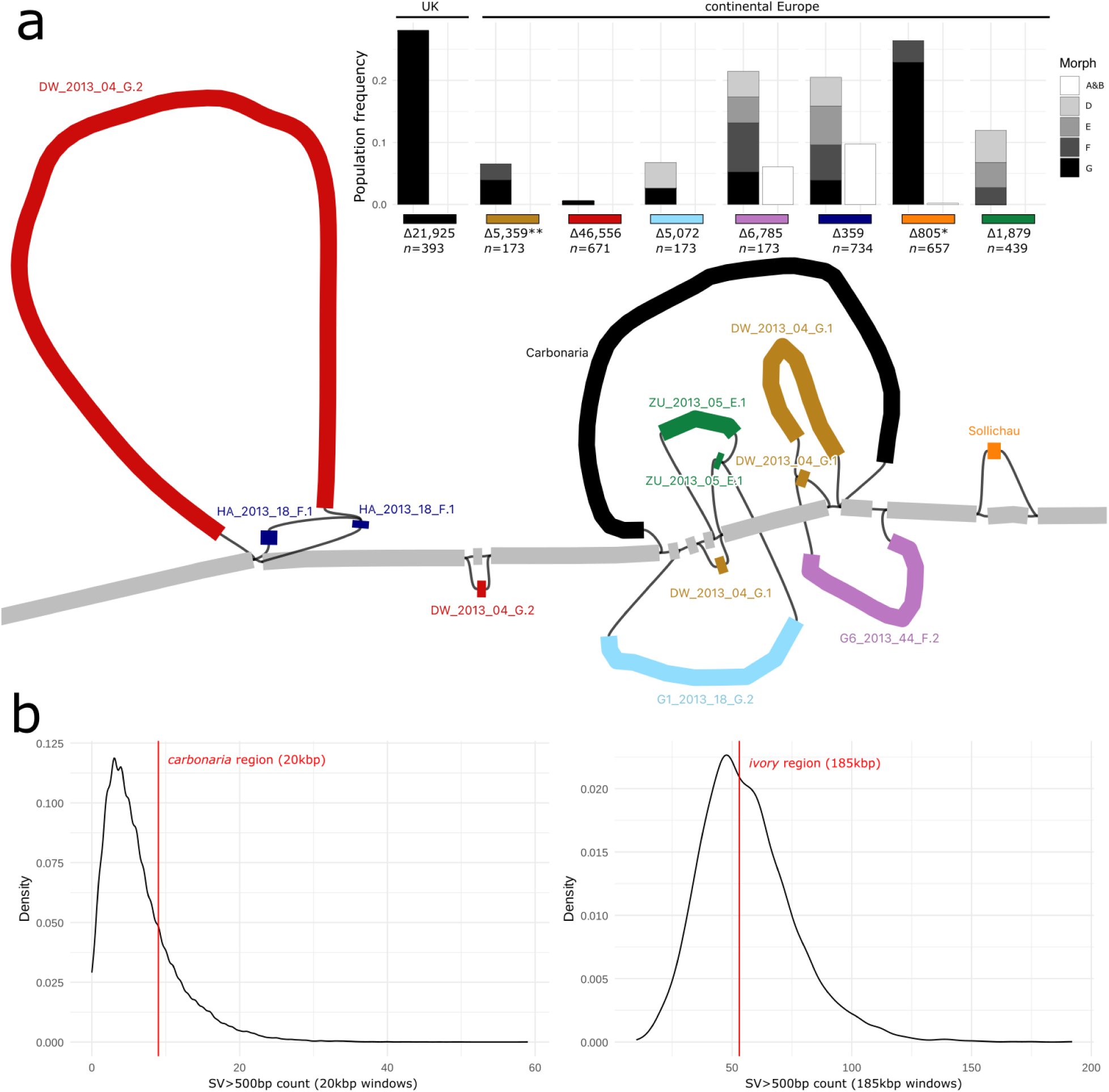
Novel structural variation found at the *carbonaria*-TE insertion site. (**a**) A pangenome graph representation of the *carbonaria*-TE and the continental European structural variants identified here. Colors represent individual sequenced haplotypes; labels specify location and year of capture, sample number, morph and haplotype 1 or 2. The sequence paths common to most haplotypes are merged to the *typica* reference backbone and represented in grey; deviations unique to a given haplotype are represented by colored paths with lengths roughly proportional to the size of the structural variant. The subplot shows the frequencies of these variants, their exact sizes in bp and their occurrence in melanic vs non-melanic morphs, among *n* individuals genotyped with short-reads. Asterisks indicate the two haplotype groups identified in Figure 4. (**b left**), The genome-wide density of structural variants > 500bp in 20kb windows across 16 haploid genomes. The 20kb window around the *carbonaria*-TE insertion is highlighted (red). (**b right**), The genome wide density of structural variants > 500bp in 185kb windows. The 185Kb window comprising the putative *ivory* ncRNA is highlighted (red).

We tested for associations of these structural variants with melanism by calling their presence/absence in our sample of 791 wild individuals sequenced with short reads using pangenome genotyping and testing for an association with melanic phenotype (AB vs DEFG). Because this region has multiple overlapping structural variants, we had varying success in genotyping. For three polymorphisms (G6_2013_44_F.2, DW_2013_04_G.1, G1_2013_18_G.2), no significant association with melanism was uncovered, possibly as we were only able to confidently call genotypes in a minority of the sample (*n* = 173). For three others (Sollichau, ZU_2013_05_E.1, HA_2013_18_F.1), a high proportion of the sample could be called (*n* = 657, 439, and 734), and these were significantly associated with melanism (*p*<0.005). When corrected for multiple testing across all genome-wide polymorphisms, only the *sollichau* deletion remains significant (Benjamini-Hochberg adjusted *p* = 1.89e-5).

### Increased *mir-193* expression is associated with melanic phenotype

We measured expression levels of *ivory* and *mir-193* in wing-disc tissues at different stages of pupal development (Figure S9) and in the context of known melanic G-morph haplotypes (*carbonaria*-insertion homozygotes and heterozygotes, and *sollichau*-deletion heterozygotes; Figure 6a; Table S6). The results varied greatly between developmental stages, with significantly higher expression of *mir-193* and *ivory* in stage 2 pupal wing discs of darker forms, and none in stage 5 wing discs. This may reflect a dynamic expression profile through time, with activity at earlier timepoints initiating pattern/color formation. Comparison among melanic genotypes does not show a straightforward linear relationship between haplotype dosage (homozygotes vs heterozygotes) and expression levels.

**Figure 6.**
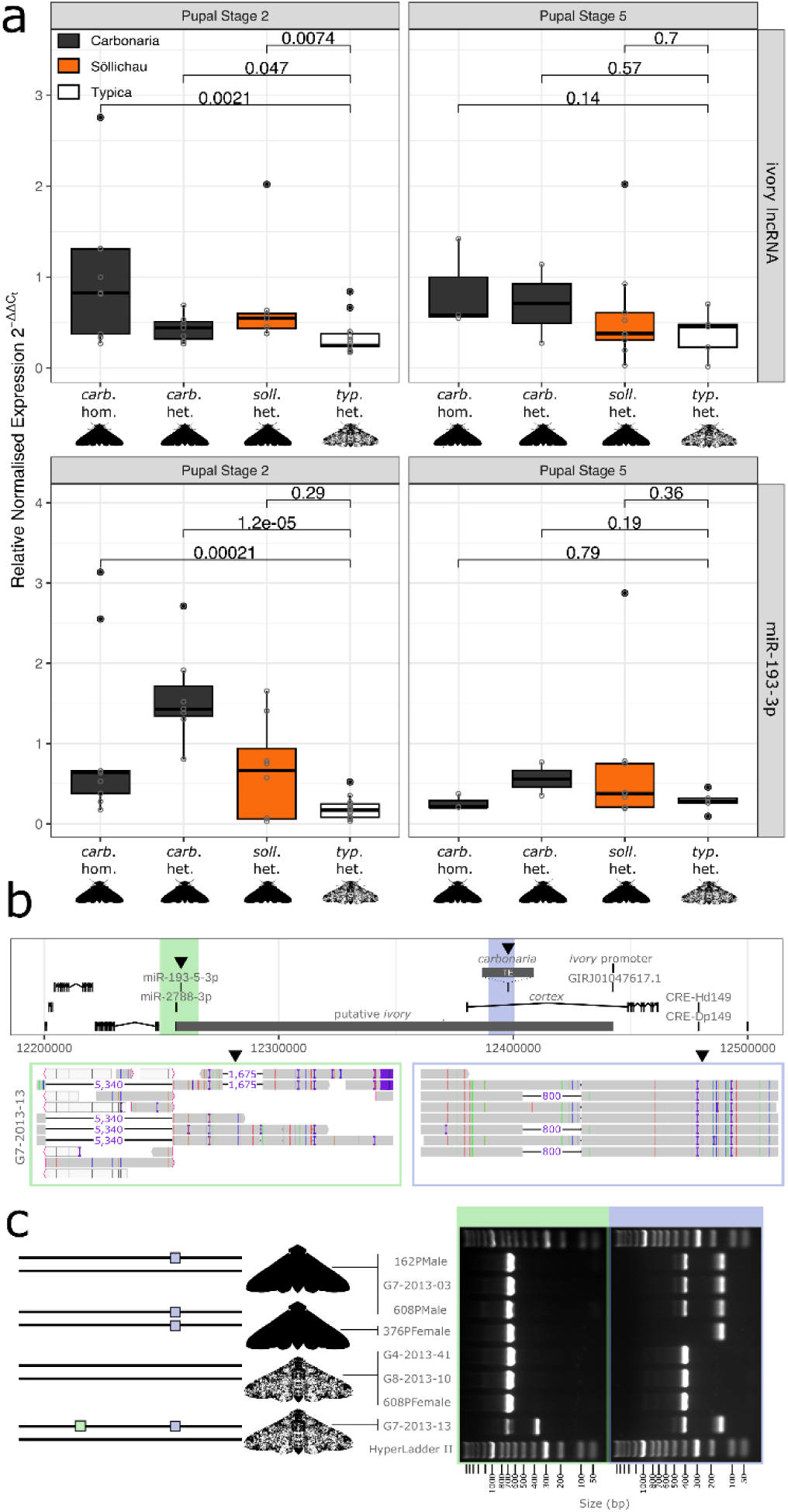
*mir-193* regulation determines melanic phenotypes and deletion of this sequence rescues a *typica* phenotype. (**a**) Relative expression levels of the *ivory* lncRNA and *mir-193* in melanic (*carbonaria* and *sollichau*) and *typica* wing-disc tissue. Wilcoxon pairwise *p*-values are indicated (range of biological *n* per genotype: 10-17 for stage 2 and 2-10 for stage 5). (**b**) Location of *mir-193* (green) relative to the *carbonaria* and *sollichau* melanic alleles (blue). Zoomed panels show a 1,675bp deletion mutant at *mir-193* (green box), discovered in a single *typica* individual carrying a *sollichau* haplotype, and the *sollichau* deletion mutant itself relative to the *carbonaria* insertion site (blue box). (**c**) A diagram showing PCR verification of the two deletions in lab-reared material and the wild *typica* individual G7-2013-13 (photo in Data S1). Phasing of the two mutations in this individual were confirmed with PacBio HiFi reads.

### A natural deletion of *mir-193* on a melanic haplotype recovers the *typica* phenotype

The candidate polymorphism for the most common melanism allele in Central and Eastern regions, the *sollichau* deletion, was present in 43 melanics (G and F) but also in one *typica* (G7-2013-13) from the Central region, raising doubts about its melanism function. This finding was supported by short-read data, PCR (on independently extracted DNA), and long-read HiFi sequencing. The long-reads further revealed that this *typica* individual was also heterozygous for a 1,675bp deletion containing *mir-193*, and that the *mir-193* and *sollichau* deletions occur on the same haplotype (Figure 6b, c). The *mir-193* deletion was not detected in any other individual within the relevant genomic region (Figure S10). We propose that, by eliminating *mir-193*, the 1,675bp deletion completely suppresses the melanisation function of the *sollichau* deletion via its cis-acting effect on increased *ivory* and *mir-193* expression. The rarity of this haplotype in the population is consistent with the expectation that lack of *mir-193* is deleterious in the homozygous state, through pleiotropic effects (weak flight) ^9,28^. Individual G7-2013-13 had normal viability and *typica* phenotype (photo in Data S1) because its alternate haplotype carried a functioning *mir-193* and lacked the *sollichau* deletion.

## Discussion

Industrial melanism in the peppered moth has served as an important case study of contemporary natural selection in response to an anthropogenic environmental change ^29^. Here, we show that the melanic allele that swept through 19^th^ century industrial Britain (*carb*-TE) did not crossover into continental Europe. Instead, several alleles appear to underlie industrial melanism on the continent, dominated by one variant in Belgium, Netherlands and western Germany, and another (*sollichau*) in eastern Germany, Czech Republic and Poland. Our analysis of museum samples from Brussels and Netherlands suggests that, even at the height of industrial melanism in the early 1900s, the darkest two morph categories (F and G) were each associated with two or three alleles. This polyallelic soft sweep underlying melanism in continental Europe contrasts the simple hard sweep of a single melanic allele observed in Britain^5^.

Despite the complexity of the underlying genetics in continental Europe, alleles associated with melanism occur in the same *ivory/cortex* genomic region as in the UK, as shown by high levels of genetic differentiation between typical and melanic morphs at this locus. In the UK, the primary casual variant is the insertion of a large (21kb) TE ^5^. In Europe, there are several structural variants in this region with high frequencies in melanic morphs. While the apparent polyallelic basis of melanism in this region reduces statistical power to detect associations, we nevertheless uncover a strong candidate for the primary G-morph allele that spread across Eastern/Central Europe, the ∼800bp *sollichau* deletion. The single *typica* individual in our sample with *sollichau* is unexpected as melanic alleles are dominant. However, this individual also has a linked deletion ablating *mir-193*, the effector gene encoded in *ivory* ^9^, consistent with a functional role for *sollichau*, and the involvement of *mir-193*. Coincidentally, the *Heliconius melpomene* mutant phenotype that led to the discovery of *ivory* was due to an even larger (78kb) deletion ^8,28^. As in other Lepidoptera ^9,10^, melanism in *B. betularia* is associated with differential expression in the pupal stage of the lncRNA *ivory* and its product, *mir-193*, rather than the previously proposed *cortex* ^5^. Intriguingly, a large insertion (*carb*-TE) appears to have an equivalent effect on the transcriptional regulation of *ivory* as a large deletion (*sollichau*).

A soft sweep, such as we propose for melanism in Europe, would be expected to leave a characteristic molecular signature of reduced diversity at the selected locus ^30^, but we find that the soft sweep statistic, H12 (quantifying the relative frequency of the two most common haplotypes) ^31^, is not elevated at *ivory/cortex*, and only modestly elevated in the flanking regions (Figure S11). One factor complicating this analysis is that modern samples are the product of a soft sweep of melanic alleles during the industrial phase, followed by a reverse sweep of the *typica* alleles in post-industrial Europe. As this reverse sweep involved former wild-type, recessive alleles that would be expected to persist at appreciable frequencies even in a fully melanic population, it was likely very soft, and thus undetectable with H12. Further, in contrast to the UK, centers of coal pollution on the continent were more dispersed, retaining large areas of low-pollution refuges where *typica* moths would have predominated and contributed to the haplotype diversity of the reverse sweep. The H12 signal we observe in the flanking region may be a remnant of the soft sweep of melanic haplotypes, which now exist mainly as recombinants with *typica* alleles at the functional locus. An analogous signature of a reverse soft-sweep was observed in British *typica* carrying *carbonaria* haplotype flanking sequence ^5^.

Why did melanism apparently arise from a hard sweep in Britain, and a complex soft sweep in continental Europe? One potential explanation is that it is an artefact of differences in the timing (and geographic coverage) of our British vs continental European samples with respect to the onset of reverse selection — which could affect the diversity of melanic alleles that were sampled. However, this explanation does not stand up to scrutiny because at the regional level the samples are comparable, both in terms of spatial scale and temporal range, even though heavy of coal pollution in Britain ended over two decades before our ‘Central’ and ‘Eastern’ European study regions (late 1960s vs 1990s). Moreover, our genetic analysis of a geographically broad sample ^32^ and the historical record in Britain are consistent with the spread of melanism from a single geographic focus (Manchester). Additional origins of melanism are unlikely to have been missed in the historical record, as amateur entomologists highly prized melanic forms in the latter half of 19^th^ century, and would have ‘assiduously’ recorded any captures, as convincingly argued by Kettlewell ^33^. In contrast, in continental Europe, melanism appears to have spread from multiple geographic foci, suggesting multiple origins of melanic alleles. A major factor that may promote soft over hard sweeps in continental Europe vs Britain is a larger effective population size (*N_e_*). In very large populations, multiple *de novo* mutations yielding the same adaptive phenotype are likely to spread simultaneously, as there are more individuals available to experience beneficial mutations. The larger spatial scale of our combined continental European sample is also likely to be a contributing factor — via barriers to the spread of melanic alleles, such as historically low levels of coal-powered heavy industry across the central German plain, promoting the spread of isolated pockets of local melanic alleles over a widespread single variant.

Equally consistent with these data, pre-industrial populations in continental Europe may have harbored standing variation for melanism in large, disconnected, areas of dark forests that seeded the multiple centers of independent industrial melanism. This hypothesis, however, is difficult to square with the convincing absence of melanic morphs in the historical record prior to localized rises in coal pollution (Table S1). Furthermore, whilst it is clear that the dominant industrial melanic in Britain was the Manchester *carbonaria*, *insularia* forms, including the F-morph, are thought to have preceded and attenuated its spread in south Wales ^33,34^, resembling the multi-allelic evolutionary dynamic we have observed on the continent. The complex, scale-dependent picture that emerges of multi-origin soft and hard sweeps within the same species is broadly consistent with theoretical predictions for a spatially extensive large population ^30,35^, and empirical studies ^10,36,37^. We have not excluded the possibility that the origin of some melanism alleles in continental Europe may precede the major environmental changes wrought by the Industrial Revolution, but their absence in the historical record and apparently large mutational target presented by the *ivory* locus suggest that *de novo* mutation during two centuries of heavy coal pollution played a central role.

## Methods

### Samples and phenotyping

Wild samples of adult *B. betularia* were principally collected using assembly trapping with large numbers of bred virgin females, supplemented with mercury vapor (MV) bulb light trapping. Descriptions of our rearing (Video S2) and assembly trapping (Video S3) procedures are available at https://dx.doi.org/10.17504/protocols.io.dm6gp92n8vzp/v1 and https://dx.doi.org/10.17504/protocols.io.8epv51odjl1b/v1, respectively. The bulk of the contemporary sample was collected in 2013 (*n* = 691), with a sample from Schevenhütte, western Germany, collected in 2014 (*n* = 47). Additional samples collected between 2000 and 2012 (*n* = 112) were used for the estimation of regional morph frequencies, with a small part of these (2007/8: *n* = 32) incorporated into the whole genome population analysis. The majority of the sample was killed and stored by freezing at −80C or −20C, with a small proportion temporarily stored in 99% ethanol.

To gain some insight into the geographic distribution of melanic alleles closer to the time period of their initial spread, we obtained tissue samples (one or two legs) from 137 melanic specimens (mostly fully melanic G morph) in three museum collections (Natuurhistorisch Museum Rotterdam, Royal Belgian Institute of Natural Sciences and Museum für Naturkunde Berlin). These specimens date from 1893 to 1981 (median year 1929), and originate largely from many locations in Brussels, Netherlands and Germany.

We scored the phenotypes according to a slight simplification of the 8-point scale described in Cook and Muggleton ^23^ into a 7-point scale (morphs A-G). As a guide, we used the photographs in Cook and Muggleton ^23^ Figure 1. The two ends of the scale (A and G) are fairly easy to score (G has no white scales on the upperside of the forewing and body) but for the intermediate morphs, as some individuals grade somewhere between the reference phenotypes, there is a degree of subjectivity in deciding which of two neighboring categories to score these as. Additionally, as specimens lose scales with age, some experience is required to distinguish wear (no scales) from white scales. To minimize these potential sources of error all of the main sample (2013/14) was scored by the same experienced researcher.

### Historical pollution and melanism data

First records of the fully melanic form of the peppered moth, *Biston betularia* L. forma *carbonaria* Jord., on the European continent were initially extracted from the early compilation of Ule ^12,13^ from which we excluded records of the intermediate melanic forms. Whenever possible, his references were traced to the original reports, leading to the correction of some of the dates (by year) of his first records. This was followed by an exhaustive literature search of further records that was facilitated by the ongoing digitization and open access to historical, multi-language journals, e.g., via the Zoological and Botanical Database ZOBODAT https://forschungsinfrastruktur.bmbwf.gv.at/en/fi/zobodat-zoologisch-botanische--datenbank_4172 and via the Biodiversity Heritage Library https://www.biodiversitylibrary.org/, which allowed cited reference searches and full text searches, using the scientific name, synonyms and spelling variations and alternative terms of the fully melanic form (e.g., *Amphidasis* Tr. *betularia* L. forma *doubledayaria* Mill.). As a result, we identified earlier accounts for 7 of the 42 locations with first records of the fully melanic form in Ule ^12^. The same location was defined as occurring within a square of 0.05 × 0.05 degrees of longitude and latitude. Our additional records covering the period investigated by Ule ^12,13^ (1867-1922) and beyond until 1963, brought the number of first occurrences recorded on the European continent to a total of 126. All records are listed with their references in Table S1).

Geographic coordinates for early records of the presence of the *carbonaria* form in Britain were extracted from Figure 1 in Steward ^34^ by overlaying his map of records onto a map of Britain. One uncertain record in south Wales (‘1860?’) was excluded.

The first occurrence records in Britain and continental Europe were used to estimate the rate of spread of the fully melanic form (some F morphs are likely to have been included in the continental European records) using a diffusion approximation approach in which the slope of the regression of distance from the very first record against year of the observation may be interpreted as the travelling wave speed of the mutation in km/generation (year) ^38^. Furthermore, the square of this wave speed is directly proportional to the relative fitness (under constant dispersal and population growth), or the axial variance in parent-offspring displacement (under constant relative fitness and population growth). This method was applied to the continental European data as a whole, under the assumption of a single geographic origin, as well as to three geographic clusters of data under the assumption of three separate geographic sources. Regressions were not forced through the origin as even highly beneficial mutants are expected to take several years to increase in frequency to the point where they would be first observed and recorded.

We retrieved estimates of black carbon particulate emissions throughout the Anthropocene by using the CEDS database ^39^. Per tile emissions within study areas were plotted through time and summed in R (v4.2.2, ggplot2 v3.5.1). We also retrieved measurements of contemporary SO_2_ levels across the 2013-2014 sampling transect from the AirBase (The European Air Quality Database, version 8, European Environment Agency) emissions dataset. The year 2010 was selected for good data coverage across continental Europe and representation of the environmental state in the generations prior to sampling. We restricted the data to rural sample locations and applied a kriging spatial-interpolation in order to estimate background SO_2_ emissions at a broad scale.

In the analysis of historical data from Czechoslovakia (1976-1982), G-morph frequencies, referred to as *carbonaria* in the original paper, were estimated from the pie charts in Figure 2 ^24^. G-morph frequencies were regressed against atmospheric SO_2_ concentration from 1995, over a decade after the sample was collected, as this was the earliest year for which AirBase records adequately cover former Eastern Bloc countries.

### Linkage mapping and pedigrees

Pedigreed lineages derived from two melanic males, referred to as Sollichau-G and Ramstedt-F, collected in 2011 from separate locations in eastern Germany (Söllichau and Ramstedt) and each crossed to bred females from the UK were maintained for several generations (original parental family identifiers were ‘162’ and ‘157’, respectively). Husbandry methods are described in Video S2 (https://www.protocols.io/edit/biston-betularia-protocols-rearing-dyj97ur6). Both of these pedigrees provided parent-offspring families used to localise the melanic alleles to chromosomal intervals via linkage mapping, and trios (father-mother-offspring) for generating long-read reference haplotypes known to carry the melanic allele. A third lineage, descended from another F-morph male collected at the Ramstedt site (‘156’) was also used for linkage mapping.

Linkage mapping was performed following the methods described in ^26^, taking into account that recombination only occurs in males in Lepidoptera. Markers fall into three categories: male informative markers (MI) are heterozygous in the father and homozygous in the mother; female informative markers (FI) are heterozygous in the mother and homozygous in the father; and both informative markers are heterozygous in both parents. MI marker genotypes can be directly used for linkage mapping, FI markers only reveal which maternal chromosome was inherited, and BI markers must be stripped of their maternal allele using FI markers to reveal the paternal allele inherited. The 156 and 157 crosses were founded by fathers which were heterozygous for a continental melanic phenotype and a mother which was homozygous typical. The 162 cross had a father which was a heterozygous continental melanic, and a UK *carbonaria* heterozygous mother. In this case, not only the BI marker genotypes, but also the phenotype had to be split into a paternal and maternal codominant component using the FI markers. The true phenotype (phenotypic morph), the excluded maternal *carbonaria* component of the phenotype (maternally inherited morph), and the paternal component of the phenotype (paternally inherited morph) are all included in (Table S3), though only the paternally inherited morph was used for positional mapping.

Primer combinations used for previous *B. betularia* studies were used to screen the fathers for segregating polymorphisms. Additionally, novel primer combinations were designed targeting polymorphisms in aligned sequences derived from *B. betularia* genome library clones (Bacterial Artificial Chromosome and fosmid) and a draft genome assembly. All these aligned sequences represent *B. betularia* from the UK, but nevertheless, heterozygous polymorphisms were also found in the fathers with a continental melanic background. The amplicons containing polymorphic indels were genotyped using band sizes on agarose gels, and SNP polymorphisms were genotyped by Sanger sequencing (Table S3). The offspring genotypes were translated to 1 and 0 depending on whether they inherited the allele from the father’s melanic or typical chromosome, respectively (Table S3).

To refine the interval further, we whole genome sequenced putative recombinant offspring identified by the marker approach. We genotyped these individuals following the same pangenome-based approach detailed below. We filtered the data for markers that were homozygous alt in known homozygous melanic individuals (Sollichau: MEL_215_F1_HOM; Ramstedt: MEL_194_F1_HOM), homozygous reference in *typica* mothers (162Pfemale [Sollichau] and 157Pfemale [Ramstedt]) and heterozygous in the long-read F1 offspring and melanic allele carrying fathers (Sollichau: 162Pmale and 181Pmale; Ramstedt: 157Pmale and 165Pmale). These markers were intersected with awk, and then phased using SHAPEIT5 ^40^ (phase_common_static), specifying the pedigree relationships. The Ramstedt sequences did not yield any meaningful improvement on the crude mapping interval.

### Preliminary study of core haplotype variation

DNA was extracted from the whole head and part thorax using either phenol-chloroform ^41^ or Qiagen DNeasy Blood and Tissue Kit. Thirty-two sequence polymorphisms within six PCR amplicons, widely spaced across 124kb were genotyped in a sub-sample of 335 individuals, by Sanger sequencing or gel electrophoresis (PCR-RFLP or indel; Table S7). Genotypes were phased with SHAPEIT2 to identify putative melanic haplogroups. Analyses of linkage disequilibrium and locus-specific *F*_ST_ profiles were uninformative, providing no clear evidence of a selective sweep driven by a single causal variant as observed in the British peppered moth. Two main hypotheses were thus considered: (i) an ancient melanic allele whose flanking regions were highly recombined; or (ii) a soft selective sweep involving several melanism alleles. We found several consistent haplogroups across sampling sites that were associated with dark morphs (E, F and G). Moreover, haplogroups associated with dark morphs did not exhibit overlapping pseudo-haplotype blocks, indicating they were not likely to derive from a common ancestral haplotype through recombination. Genotypic data therefore appeared more in line with the spread of several distinct alleles, although a single allele scenario, with complex demographic processes such as historical bottlenecks or periods of population isolation could not be completely ruled out at this stage.

### Reference haplotypes

To obtain phased melanic haplotype assemblies for Sollichau-G and Ramstedt-F alleles, we sequenced a melanic F_1_ offspring from melanic × *typica* crosses for each of these alleles with PacBio HiFi reads, and performed short-read WGS on mothers and fathers. The recorded parentage of these HiFi trios was confirmed by examining the intersections of read 31-mers (meryl v1.3 ^42^). Based on distinctness of PCR-marker profiles in the *ivory/cortex* region (and morph), we additionally sequenced 6 wild-caught melanic individuals from across the study area with PacBio HiFi reads.

Samples were taken from family material frozen immediately after eclosion and phenotype scoring, or after mating, and stored at −20C or −80C prior to extraction. High molecular weight genomic DNA was extracted using Qiagen Genomic-tips, and was used as initial material for low-input HiFi libraries constructed by the Centre for Genomic Research (Liverpool UK). Libraries were sequenced on a single PacBio Sequel II SMRT Cell 8M, with a subset of libraries requiring resequencing to increase coverage.

The PacBio reads were assembled with hifiasm (v0.15.1-r334), in either haplotype or diploid mode, depending on availability of parental data. The resulting subsequent partitioned or mosaic haplotype genomes were combined into a pangenome graph, using a reference genome as the coordinate system (NCBI accession: GCA_905404145.2), using the Minigraph-Cactus Pangenome pipeline (v2.9.0).

### Population-level WGS analyses

Whole genome sequence for modern samples was obtained by double-stranded library prep (NEBNext Ultra II FS DNA library prep kit for Illumina) with the minimum number of PCR cycles (3), followed by short-read Illumina sequencing to 15x median coverage. For museum samples, one or two legs were collected into LoBind tubes and extracted with QIAamp DNA Micro Kit. Genomic libraries for museum DNA were made using the Santa Cruz Reaction protocol ^43^, which is specifically designed for templates with a high proportion of single-stranded DNA. The amount of starting DNA for museum libraries, based on ds measurement, ranged from x-y, and they were all subjected to 8 PCR cycles. 137 museum libraries were combined into three pools, each sequenced on a lane of Illumina NovaSeq 6000, yielding a median mapped coverage of 9x. The entire process of DNA extraction and library prep for museum samples was conducted in a separate laboratory where no invertebrate or other eukaryote work was conducted.

We genotyped the samples sequenced with short-reads against the pangenome using a k-mer based inference method (PanGenie v2.1.1) and filtered by Genotype Quality greater than 200. We called SNP and small variants *de novo* from the same samples (excluding the museum individuals) using freebayes (v1.3.6) (-p 2 --use-best-n-alleles 4 -m 20). We also called larger structural variation *de novo* using dysgu (v1.7.0 --min-support 2) and included the long read samples (--mode pacbio-sequel2), including two individuals that were absent from the pangenome reference due to low coverage.

For principal component analysis of the samples across the genome we first filtered sites with a max missingness of 0.95 and minor allele frequency 0.05. For the whole genome PCA we also performed LD pruning using bcftools (v1.11 with a threshold of 0.25 and window of 1kb). Principal components were calculated and explored using pcadapt (v4.3.5).

For visualization of multilocus genotypes, genotyping files were first normalized with bcftools, to represent multiallelic sites individually. These genotype files were then tested independently for association with melanic phenotype categories (A & B individuals vs D, E, F, G individuals) using a chi-square allelic test with 1df, implemented in PLINK (v1.90b6.21). We then selected loci from each contrast to visualize multilocus genotypes significantly associated with melanism (D, E, F threshold: p<0.01). A more stringent significance threshold was used for the G group to yield a more manageable number of loci for visualizations (p<1e-8). Prior to visualization, we performed phasing and imputation of the genotype files using SHAPEIT4. We calculated and plotted a minimum spanning MLG network using poppr (v2.9.4) using a threshold of 6 to contract similar MLG groups.

To calculate the soft-sweep statistic H12 across the genome, we used the same phased genotype files as an input to LASSI-Plus (v1.2.0) with window size and step parameters 2500 and 500, respectively.

### Structural variation

Pangenome structure at the *carbonaria* locus was visualized with Bandage (v0.8.1). The genome-wide distribution of structural variants was measured in windows across the genome using bedtools (v2.31.1). For windows comparable to the *carbonaria* insertion site, we used 20kb windows in 10kb steps. For regions comparable to the *ivory* locus as a whole we used windows of 185kb in 92.5kb steps.

PCR validation was performed for selected structural variants. Flanking and internal primers were designed using primer-BLAST ^44^. Three-primer combinations were used to produce distinct amplification products in both presence and absence of a structural variant (Table S7).

Structural variants with homology to transposable element sequences were curated from the reference and pangenome sequences, and classified in accordance with the Wicker et al ^45^ classification system using a combination of diagnostic information from examination of conserved protein domains, coverage plots, and target site duplications for each sequence.

### Gain- and loss- of function deletions

PanGenie genotyping suggested that a single typica individual (G7-2013-13) carried the most strongly associated melanism SV, a deletion detected originally in the Söllichau trio. We also applied a *de novo* structural variant calling approach to the short-read data, and detected a 1,675bp deletion in individual G7-2013-13, this time deleting the candidate melanism effector sequence for *mir*-193.

To investigate further we generated additional PacBio HiFi data for individual G7-2013-13. The coverage was not high enough to assemble contiguously, however the mapped HiFi read data confirmed the presence of both the *sollichau-*deletion and the *mir-*193 deletion. Phasing the called variants with WhatsHap (v2.3), confirmed that both deletions were present on the same haplotype. We validated this further in G7-2013-13 by designing three-primer PCR assays, wherein two flanking primers and a third, deletion-specific internal primer generates different fragment lengths based on presence, absence and zygosity of the deletion polymorphism (Table S6).

### Expression of *ivory* and *mir-193*

RT-qPCR was performed to quantify the guide strand of *mir-193*, miR-193-3p, and its primary transcript, *ivory*, among alternative melanic (homozygous and heterozygous) and non-melanic (homozygous) genotypes within a small number of sibships. Melanic alleles were represented by *carbonaria*-TE (homozygous and heterozygous state) and s*ollichau*-deletion (heterozygous state only). Pupae were genotyped using 3-primer PCR, diagnostic for *carbonaria*-TE ^5^ or s*ollichau*-deletion (Table S7). Fresh pupal wing discs were dissected at post-diapause stages 2 and 5 (Figure S9). All four wings from one individual were pooled as one biological replicate. Data were preferentially collected for stage 2 over stage 5, such that 10-17 biological replicates per gene were obtained for stage 2 and 2-10 for stage 5. Fresh tissues were preserved in RNAlater before RNA extraction. Owing to variation among individual pupae in their response to incubation temperature profiles, it is not possible to accurately determine post-diapause developmental stage prior to removal of the cuticle. Wing disc developmental stage was therefore determined based on their appearance (Figure S9).

Total RNAs, including small RNAs, were extracted using miRNeasy Tissue/Cells Advanced Mini Kit (Qiagen). To quantify miR-193-3p, cDNAs were reverse-transcribed from 1μg total RNAs using a stem-loop (SL) primer, together with another SL primer for the small nuclear RNA (snRNA) U6 as an internal control ^9^. To quantify *ivory*, cDNAs were reverse-transcribed from 1μg total RNAs using oligo-dT. *RpS3A* was used as an internal control ^5^. Reverse transcription was performed using RevertAid RT Reverse Transcription Kit (Thermo Fisher). To quantify miR-193-3p, cDNA products from the reverse transcription reaction were used for qPCR using a miRNA-specific qPCR forward primer, and a universal qPCR reverse primer ^9^. To quantify *ivory*, qPCR primers were designed to amplify a 80bp region from the ivory first exon, close to the deeply conserved *ivory* transcription start site (TSS). qPCR was performed using KAPA SYBR FAST Universal Kit (Roche). The qPCR thermal cycling program was set as 40 cycles of 95°C for 10s and 60°C for 30s. For each biological replicate, three technical replicates were included, and expression data was analysed using the 2^-ΔΔCt^ method. Melting curves were generated after each qPCR run and checked to ensure unique amplification. All SL primers and qPCR primers are listed in Table S7.

## Data availability

Sequencing data and reference genome assemblies are available at ENA BioProject accession PRJEB95936 (from 01/01/2026). The sequences of *ivory/cortex* region structural variant insertions putatively associated with melanism are provided in Supplementary Data 2. Other data supporting the findings presented in this paper are available in the Supplementary Tables. Photographs of a large proportion of the wild-caught peppered moths analysed for this study are available at the Zenodo repository https://doi.org/10.5281/zenodo.14945781.

## Acknowledgements

We thank the following people for collecting or facilitating the collection of contemporary *Biston betularia* material: Paul Brakefield, Christian Shulze, Otto Elias, Josef Settele, Hynek Habal, Henk Spijken, Jan Heesters, Bas Zwaan and Karin Geertsen. The genetic analysis of historical samples was possible thanks to the permission and assistance of curators at three museums: Royal Belgian Institute of Natural Sciences (Wouter Dekoninck and Stefan Kerkhof); Museum für Naturkunde Berlin (Théo Léger and Jannes Stickamp); and Natuurhistorisch Museum Rotterdam (Kees Moeliker). Ivo Novak and Karol Spitzer kindly shared unpublished records of melanism frequency in Bohemia. Amy Corthine, Rosie Griffiths, Fatema Khambati, Daisy Collins, Becky Court, Scarlett Zetter, Eve Taylor-Cox and Kieran Romain assisted with husbandry and sample processing. Josh Kapp helped us to troubleshoot the Santa Cruz Reaction genomic library prep. Centre for Genomic Research (Liverpool) staff provided expert support in sequencing (Charlotte Nelson, Margaret Hughes, Anita Lucaci, Ecaterina Vamos and Chris Owen) and bioinformatics infrastructure (Richard Gregory). This research was primarily funded by Natural Environment Research Grants (NE/J022993/1 and NE/T000597/1). We also acknowledge the National Research Foundation Singapore, Competitive Research Program (award NRF-CRP20-2017-0001).

## Supplementary figures

**Figure S1.**
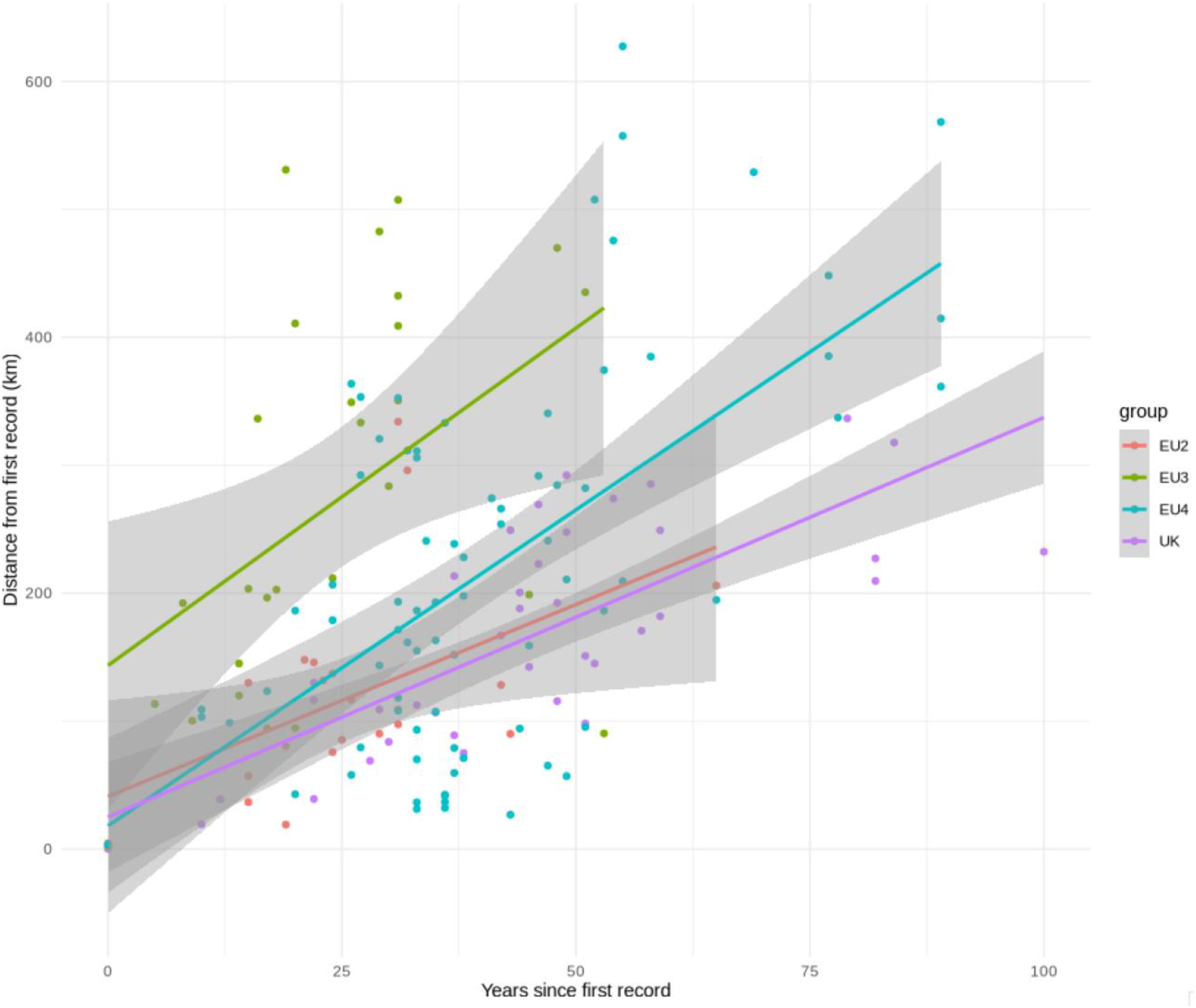
An alternative multi-origin model for the spread of melanic *Biston betularia* (G and some F morphs) in western Europe. Linear regressions were fitted to records within the four main geographic clusters shown in Figure 1a, putatively spreading from first occurrences in 1848 (UK), 1867 (EU2), 1879 (EU3) and 1874 (EU4). The slopes, representing the rate of spread in km/year, of the EU clusters [3.0 (EU2); 5.3 (EU3); 4.9 (EU4)] are more similar to the UK (3.1) than the EU single origin model (Figure 1c), though still noticeably higher for EU3 and EU4. The shaded areas indicate the 0.95 confidence intervals for each slope.

**Figure S2.**
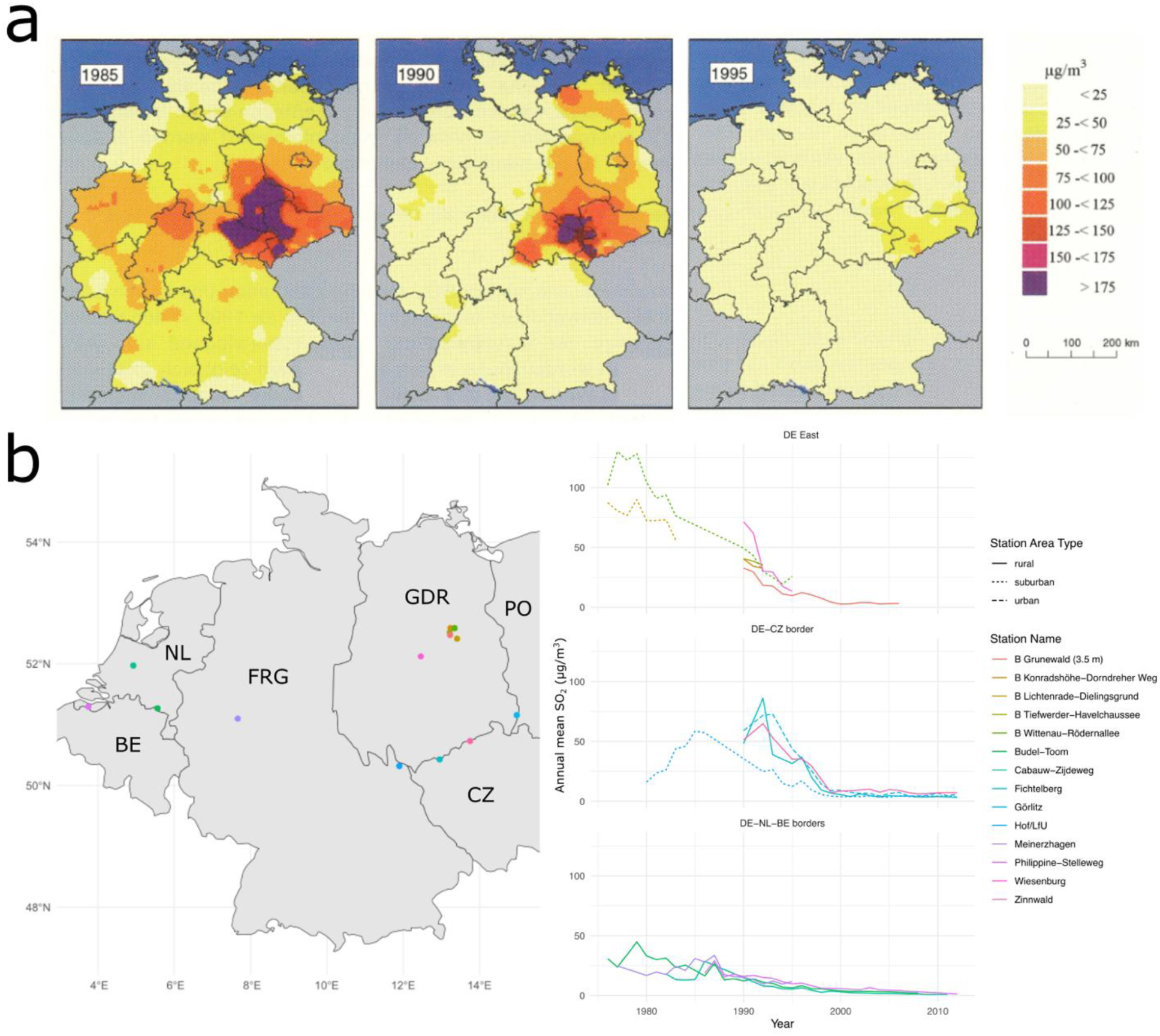
Spatio-temporal patterns of SO_2_ concentration in Germany and neighboring countries with extensive industrial and domestic coal use. (**a**) Contrasting SO_2_ concentrations between the former Federal Republic of Germany (FRG, West Germany) and German Democratic Republic (GDR, East Germany), reflecting the main coal extraction areas, much earlier introduction of atmospheric pollution controls in West Germany, and the rapid elimination of the SO_2_ gradient following German reunification in 1990 ^46^. (**b**) Graphs showing the overall decline in annual mean SO_2_ concentrations between 1976 and 2013, highlighting the much higher concentrations in the Central and Eastern study regions (DE East and DE-CZ border) compared to the Western study region (DE-NL-BE borders), and the sharp decline from 1990 in the Central and Eastern regions. The locations of the SO_2_ measuring stations are shown in the outline map of Germany (divided into FRG and GDR), Belgium (BE), Netherlands (NL), Czech Republic (CZ) and Poland (PL). Station names prefixed with ‘B’ are in the Berlin area. AirBase data from some recording stations are incomplete, and very limited before 1990 for Central and Eastern regions.

**Figure S3.**
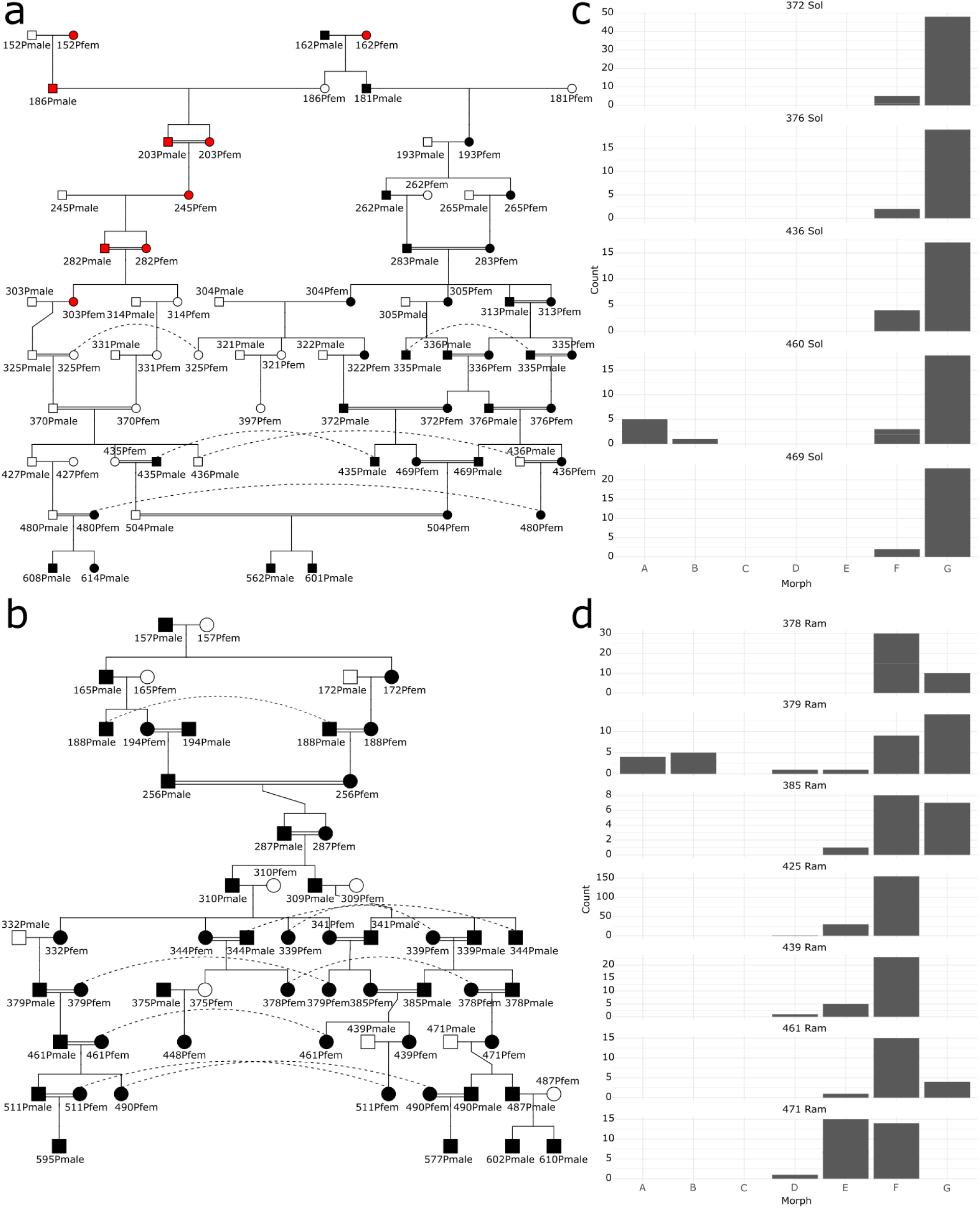
Pruned pedigrees of the Sollichau-G (**a**) and Ramstedt-F (**b**) melanic alleles used for recombination mapping. Photographs of the original wild-caught male parents (162Pmale and 157Pmale) and one bred female parent (162Pfem) are shown in Figure S5 (black symbols - melanic phenotype; white symbols - *typica* phenotype; red symbols - UK *carbonaria*-TE). (**c**, **d**) Phenotype distributions in the offspring of a sample of Sollichau-G and Ramstedt-F families represented in the pedigrees.

**Figure S4.**
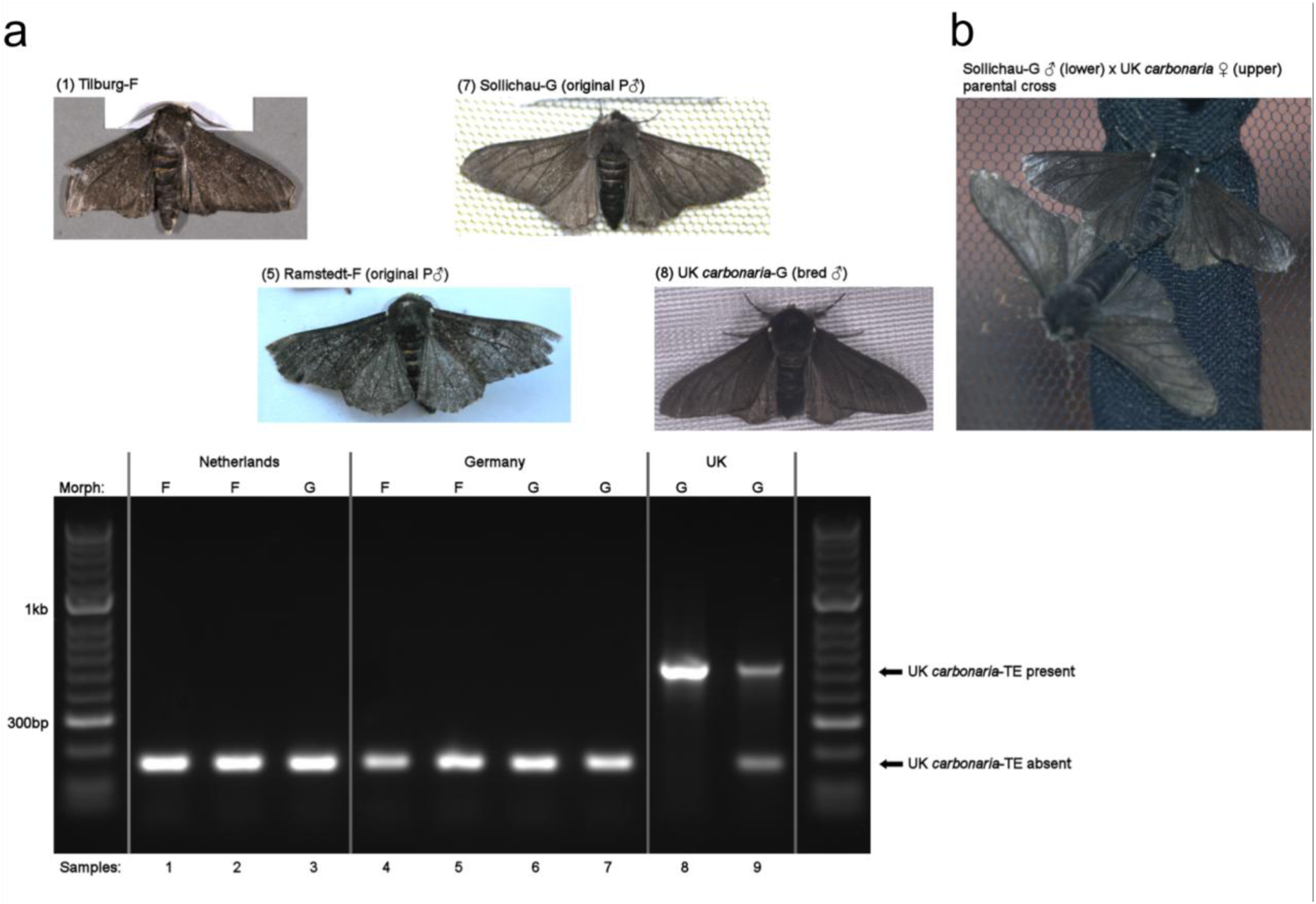
(**a**) Initial evidence for the absence of UK *carbonaria*-TE in continental European melanics. Diagnostic 3-primer PCR assay ^5^ in a sample of F and G morph melanics from Netherlands (2007 and 2008), Germany (2011), and UK (2002 and 2016). As the *carb*-TE allele is dominant, homozygotes (sample 8) and heterozygotes (sample 9) express the same melanic (G) phenotype. Images of some of the genotyped moths, including the original wild-caught fathers for the Sollichau-G and Ramstedt-F alleles, are linked to their corresponding UK *carb*-TE genotype by sample numbers. (**b**) Original parental cross of the Sollichau-G allele. The female in the cross was UK *carbonaria* (heterozygous for *carb*-TE and *typica*) rather than the preferred homozygous *typica* as no other material was available.

**Figure S5.**
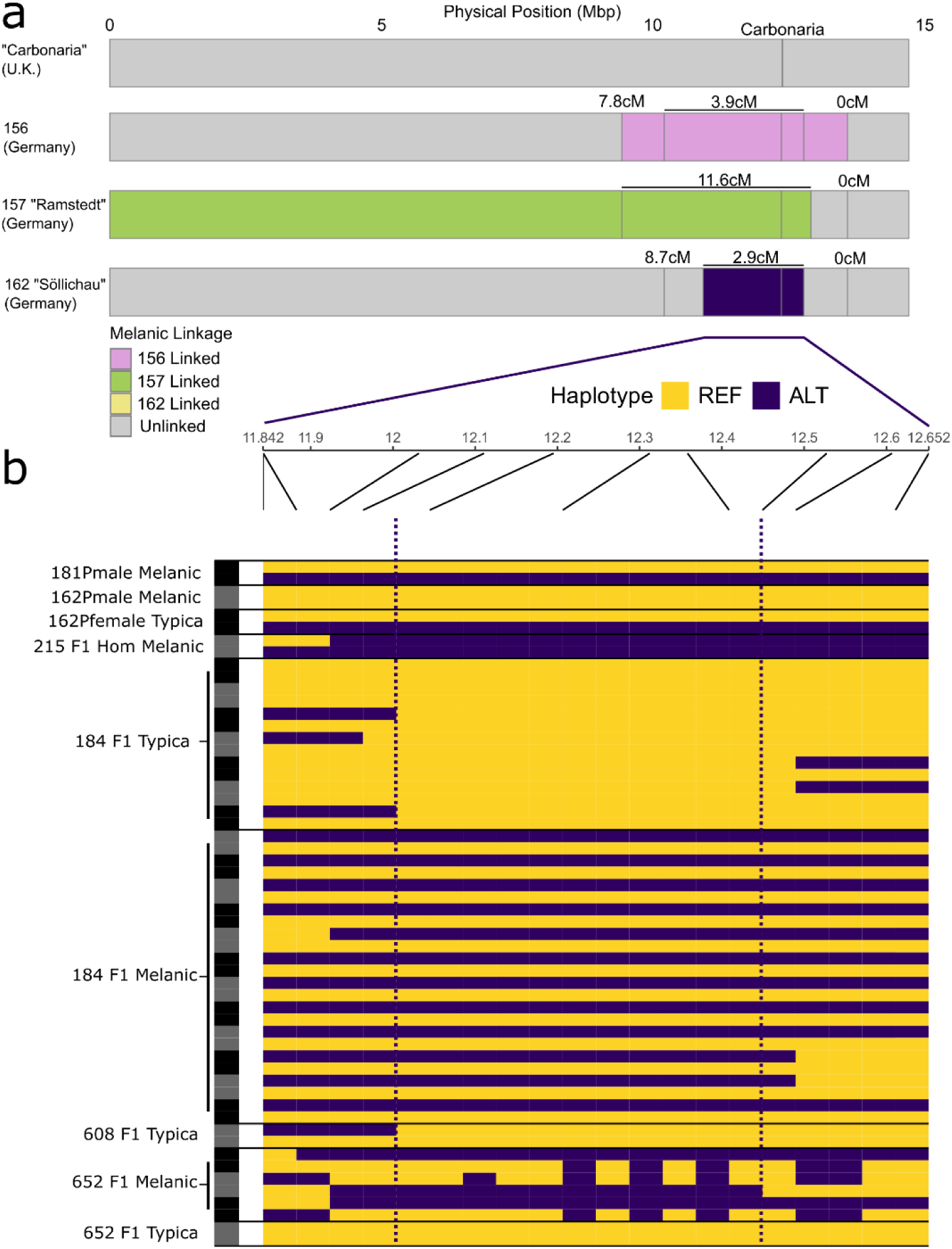
Linkage mapping intervals for three continental European melanism alleles. (**a**) The physical position (with annotated genetic position in cM) of gene markers used to generate coarse linkage maps of three European alleles: Sollichau-G; Ramstedt-F; and ‘156’, another F-morph caught at the Ramstedt site (numbers are family identifiers). *Carbonaria*-TE position is shown for reference. (**b**) A fine-scale (Mb) display of phased genotypes in a sample of descendants of Sollichau-G (parental 162 cross shown in Figure S3a and S4b), showing precise recombination breakpoints (melanic associated variants shown in dark blue). Individuals are grouped by F1 sibships and phenotype (none of these families are shown on the Figure S3a pedigree). Dotted lines define the 417Kb interval (Chromosome FR989875.1:12,110,822 – 12,527,387) for the Sollichau-G allele.

**Figure S6.**
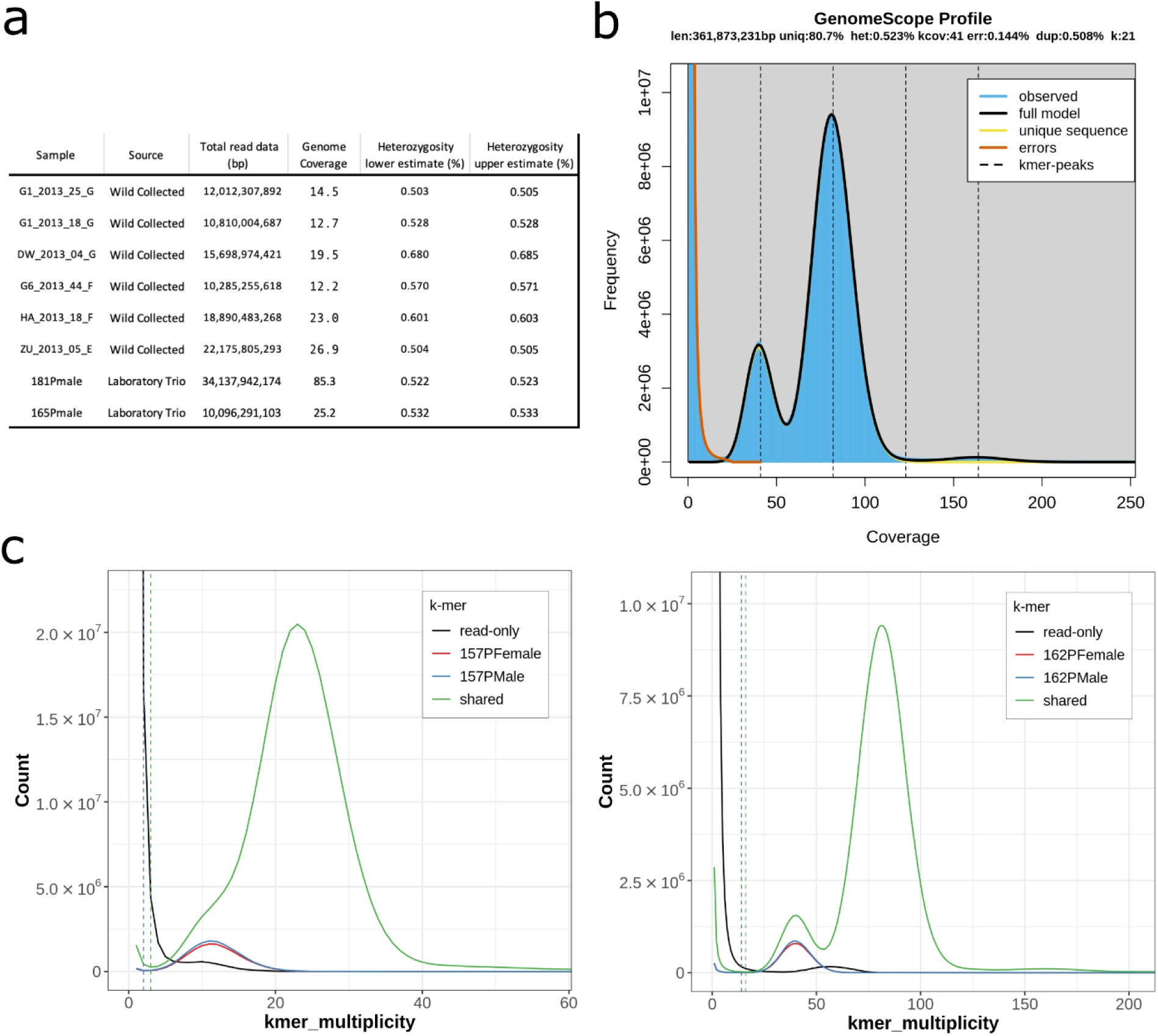
PacBio HiFi data QC utilising k-mer spectra. (**a**) Table describing overall sequencing output, coverage and heterozygosity estimates. (**b**) GenomeScope k-mer histogram plot for the highest coverage individual (181Pmale) and the associated fitted content estimation models. (**c**) Shared k-mer content between parents and offspring for the two laboratory cross trio individuals, left “Ramstedt” (165Pmale) and right “Söllichau” (181Pmale).

**Figure S7.**
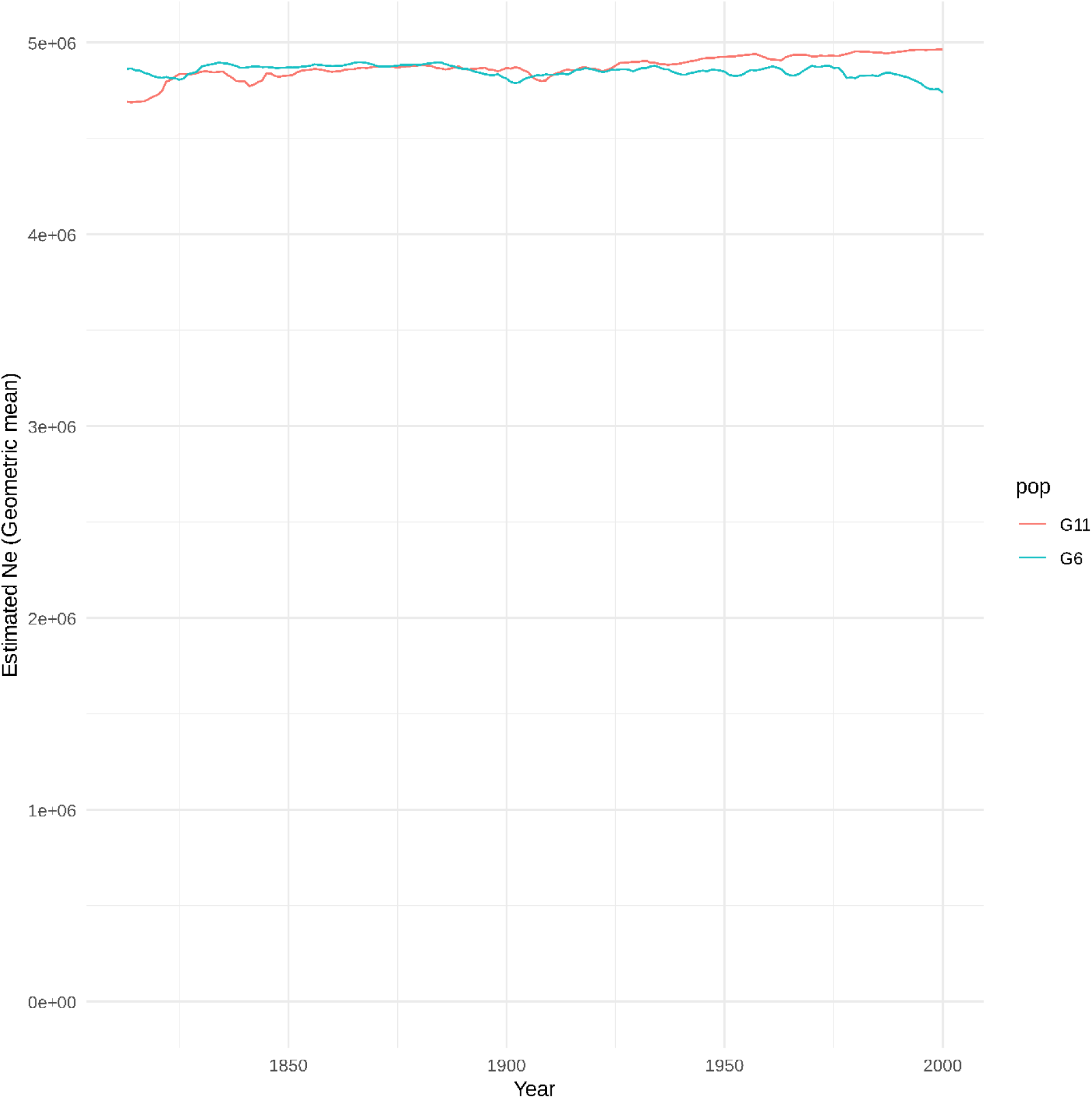
Effective population size (*N*_e_) of *Biston betularia* over the past 200 years (generations) across the study region in continental Europe. The estimated profiles were inferred from the pattern of linkage disequilibrium ^47^ in two population samples collected in 2013/14 (blue - G6 Deutzen in eastern Germany, *n* = 38; red - G11 Schevenhütte in western Germany, *n* = 43).

**Figure S8.**
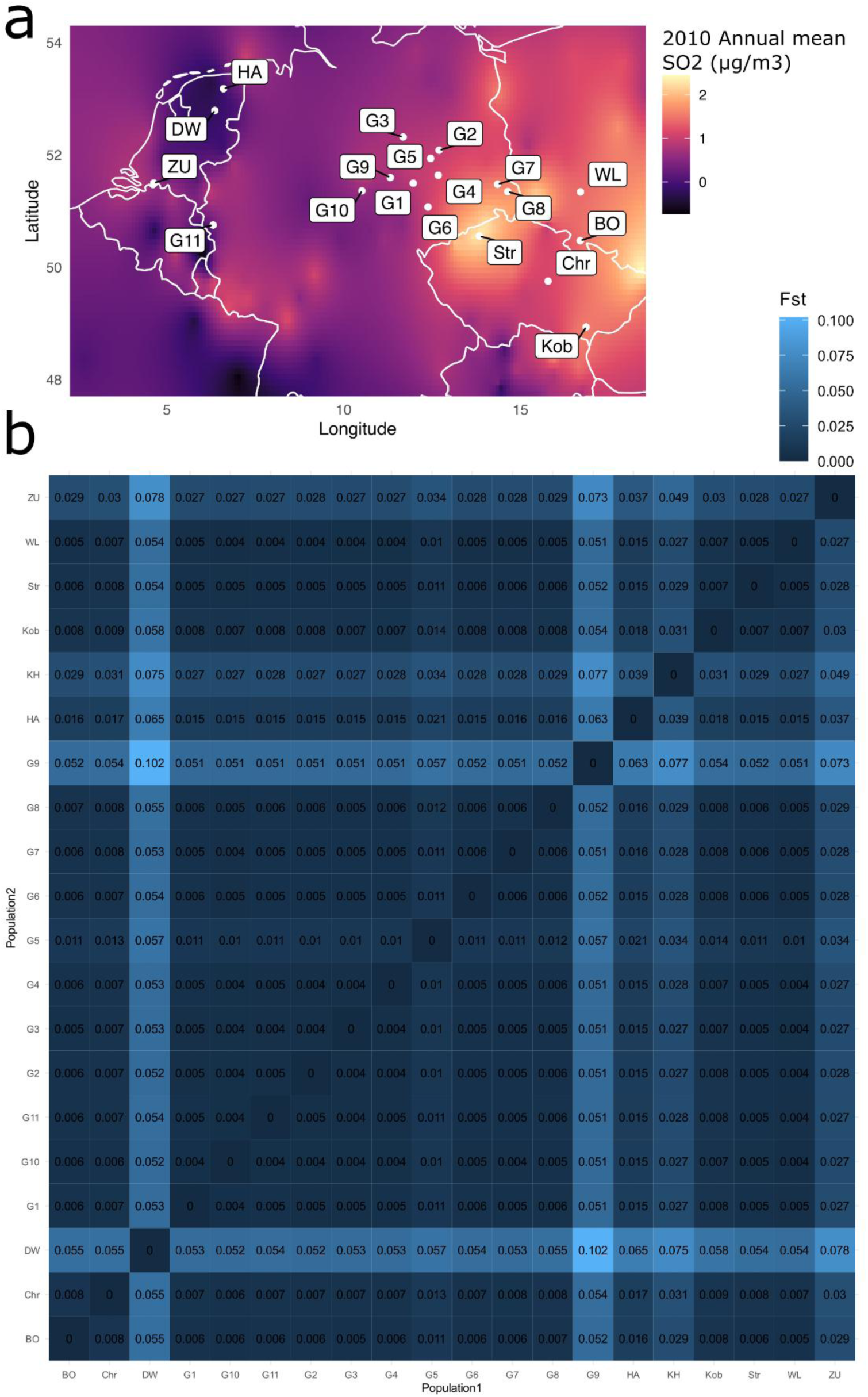
(**a**) Outline map of north-western continental Europe showing the locations of *Biston betularia* samples collected in 2013 and 2014, in the context of atmospheric sulfur dioxide concentration in 2010. (**b**) Matrix of genome-wide pairwise *F*_ST_ between sampling locations.

**Figure S9.**
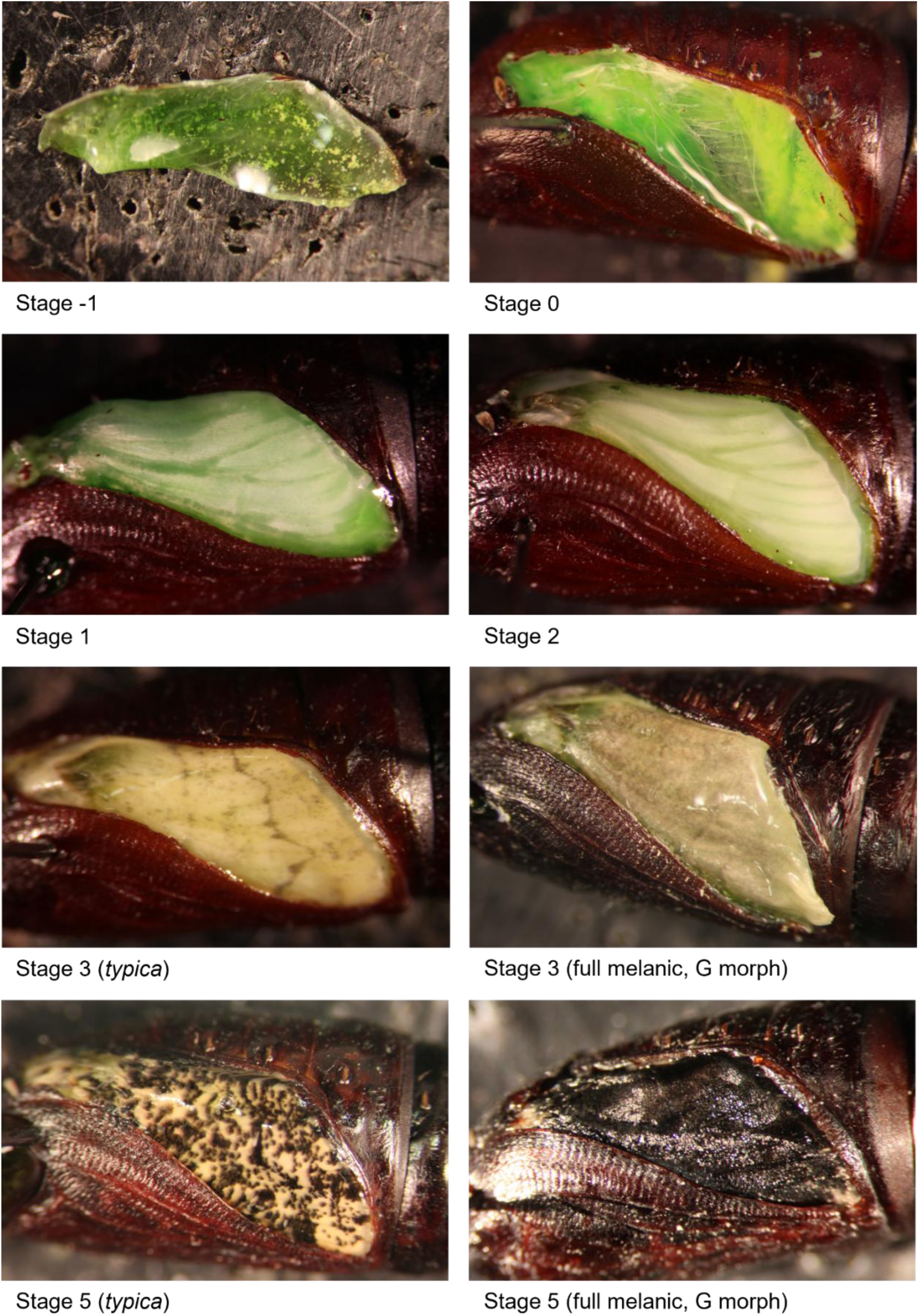
Developmental stages of *Biston betularia* wings in post-diapause pupae (cuticle partially removed). At stage −1, undeveloped wing tissue is stuck to the pupal cuticle; at stage 0, the wing remains undeveloped but is detached from the cuticle. Visible pigmentation, and therefore differentiation among forms, begins at stage 3 and is complete by stage 5. Adult eclosion follows stage 5 (stage 4 not shown). At 20°C, development from stage 0 to 5 takes 7-8 days.

**Figure S10.**
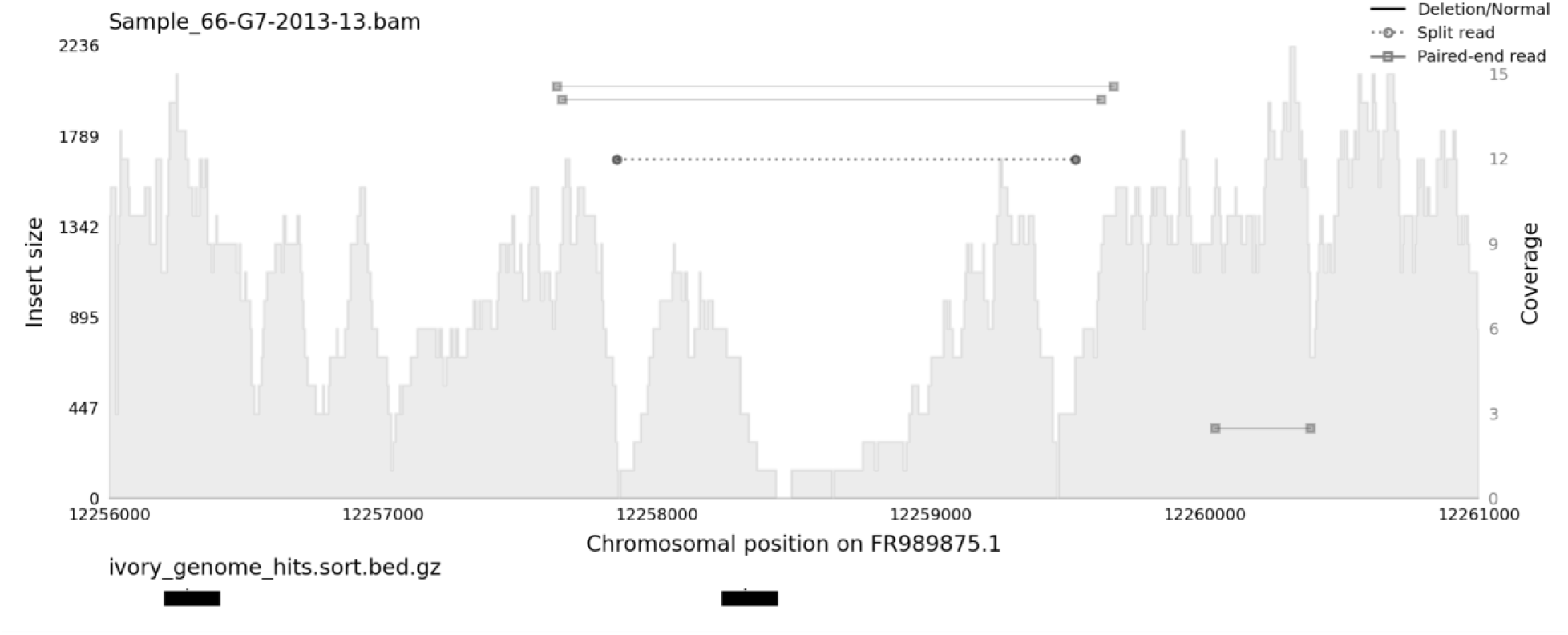
Samplot image showing coverage, insert size, and split reads at the 1,675bp deletion mutant identified in G7-2013-13. A shaded histogram indicates the coverage across the region (right-hand axis). In addition, individual read pairs or split reads with unusually large insert sizes are displayed as an overlay (left hand axis). Together, these signals corroborate the presence of a deletion at *mir-193*.

**Figure S11.**
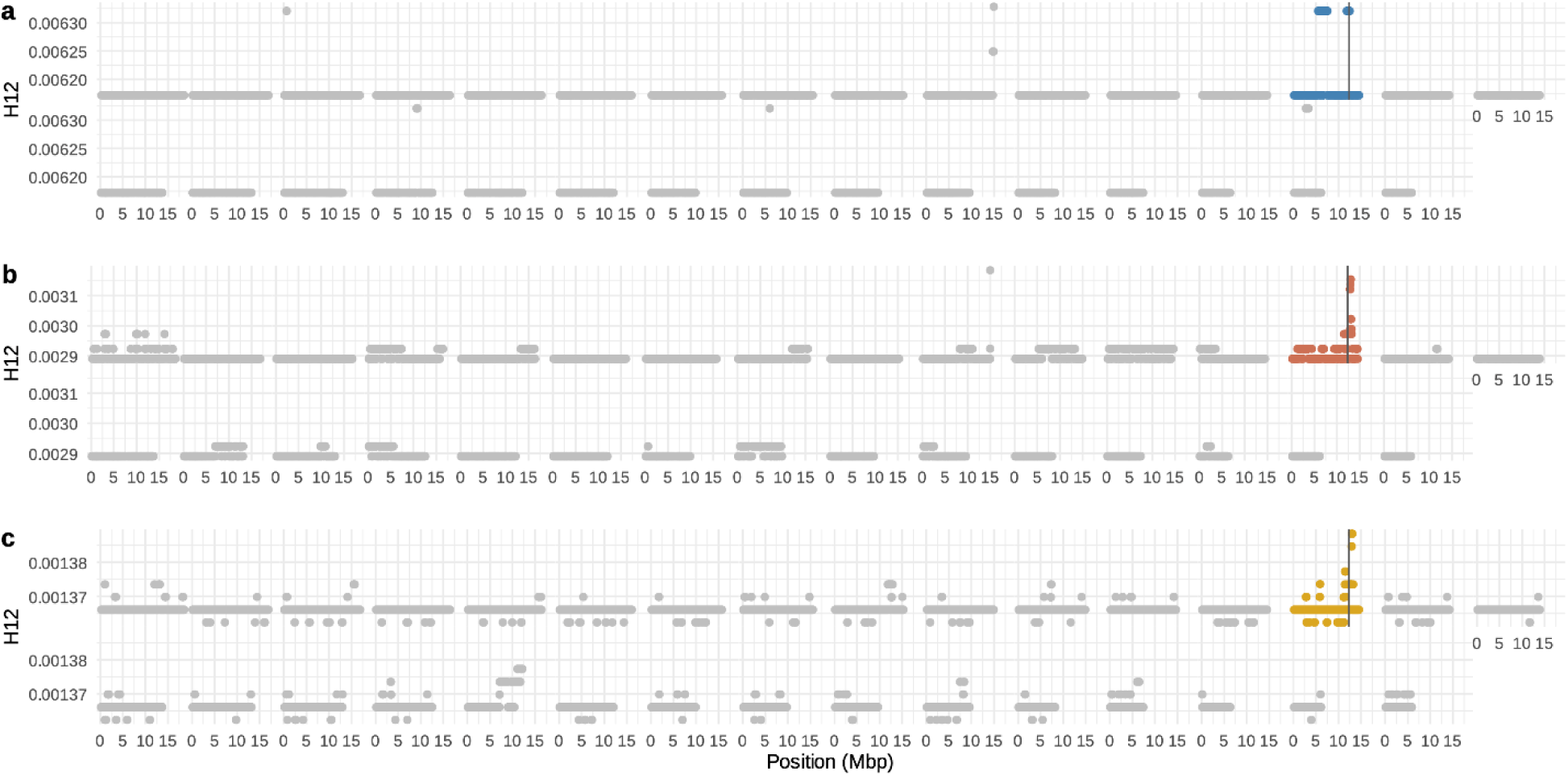
Genome-wide windows (size 2500, step 500) of the soft sweep statistic H12 in modern sample, broken down by the three study regions: Western (**a**), Central (**b**) and Eastern (**c**). The chromosome containing ivory is coloured, whereas values across other chromosomes are gray. Y-axes scales are variable between populations due to differing sample size and reflecting different selection regimes, emphasising the position rather than the magnitude of the signal. The vertical black lines indicate the Sollichau-G mapping interval.

## Supplementary information

Table S1. Excel spreadsheet of early records of melanic *Biston betularia* for continental Europe and Britain, together with historical background text.

Table S2. Excel spreadsheet of metadata associated with all sequenced individuals collected from the wild (modern – 2007-2014 and museum – 1893-1981).

Table S3. Excel spreadsheet of genotype matrices for mapping families (and marker details), and metadata for pedigreed individuals that were sequenced (for trio assemblies and recombinant mapping).

Table S4. Excel spreadsheet of PCR-marker *ivory/cortex* region phased genotypes, analysed in a preliminary study.

Table S5. Features and classification of structural variants in the *ivory/cortex* region found to be potentially associated with melanism.

Table S6. Cycle Threshold (Ct) values obtained by a RT-qPCR experiment to compare relative expression levels of long non-coding RNA *ivory* and microRNA *mir-193* in developing wing discs of melanic and non-melanic morphs of *Biston betularia*.

Table S7. Various PCR primers: *ivory/cortex* region exploratory study; 3-primer PCR (*sollichau* deletion, *mir-193* deletion, *carb-TE*); RT-qPCR.

Video S1. Animation of progress of SO_2_ concentration, a correlate of soot pollution, in northwestern and central Europe, from 1750 to 2019. Data extracted from the Community Earth atmospheric Data System (CEDS).

Video S2. Protocols.io *Biston betularia* husbandry methods (DOI: dx.doi.org/10.17504/protocols.io.dm6gp92n8vzp/v1).

Video S3. Protocols.io *Biston betularia* trapping methods (DOI: dx.doi.org/10.17504/protocols.io.8epv51odjl1b/v1).

Data S1. Photographic database of some of the samples listed in Table S2, including the double deletion (*sollichau* and *mir-193*) *typica* individual G7-2013-13 and three melanics (G) from 1946 (https://doi.org/10.5281/zenodo.14945781).

Data S2. FASTA file of the structural variant insertion sequences detected in the *ivory/cortex* region that are putatively associated with melanism, described in Table S5

